# GABAergic interneuron pathology in schizophrenia: a systematic review and meta-analysis across cell-types, brain areas, and cortical layers

**DOI:** 10.1101/2025.05.23.655812

**Authors:** Aidan G. Mulvey, Kaitlyn M. Gabhart, Tineke Grent-’t-Jong, Suzana Herculano-Houzel, Peter J. Uhlhaas, André M. Bastos

**Affiliations:** Department of Psychology, Vanderbilt University, Nashville, TN; Department of Psychology, Yale University, New Haven, CT; Department of Clinical and Counseling Psychology, Columbia University, New York, NY; Department of Child and Adolescent Psychiatry, Charité-Universitätsmedizin, Berlin; Vanderbilt Brain Institute, Vanderbilt University, Nashville, TN

## Abstract

**Introduction:** GABAergic interneurons are implicated in the pathophysiology of schizophrenia, yet evidence regarding the nature of deficits across brain areas and interneuron subtypes remains conflicting. We adopted a meta-analytic, multi-level linear modeling approach to identify interneurons, cortical layers, and brain areas involved, and the implications for circuit functions in schizophrenia.

**Methods:** Following PRISMA guidelines, we conducted a systematic search and meta-analysis from inception to November 2025, for studies examining parvalbumin, somatostatin, calbindin, and calretinin interneuron density or mRNA expression in schizophrenia. We included data from 44 studies, comprising 736 individuals with schizophrenia and 814 healthy controls. Non-cell-specific, non-human, or indirect proxy studies were excluded. Linear mixed-effects models quantified deficits while accounting for cortical layer, cell-type, and brain area, providing a map of interneuron pathology. We further analyzed changes in GABAergic interneurons to determine whether deficits preferentially target cortical layers, cell-types, and brain areas more associated with *top-down* or *bottom-up* processing.

**Results:** Parvalbumin and somatostatin interneurons showed robust reductions, particularly in layers 3/4, whilst calbindin and calretinin interneurons were less affected. Deficits were widespread across cortical and subcortical regions. Contrasts revealed that schizophrenia is characterized by interneuron deficits preferentially affecting bottom-up signaling—notably parvalbumin and somatostatin interneurons in layers 3/4, which are critical for gamma-band synchronization and feedforward sensory processing.

**Discussion:** These findings provide the most comprehensive meta-analysis on GABAergic interneurons in schizophrenia to-date, as well as a novel perspective on circuit dysfunctions, with implications for computational models. These results highlight the need for more widespread sampling across the brain using methodologies that can pinpoint deficits in molecularly more precise ways.

## INTRODUCTION

Schizophrenia is a severe psychiatric disorder characterized by positive symptoms, such as delusions and hallucinations, negative symptoms, and cognitive deficits [1]. Aberrant dopaminergic [2], glutamatergic [3], and GABAergic [4,5] neurotransmission has been proposed in circuit models of schizophrenia [1], yet the underlying mechanisms and localization of circuit deficits remain unclear.

Several cortical and subcortical areas, including the prefrontal cortex (PFC) [6], hippocampus [7], thalamus [8], and other subcortical loci [9] have been implicated in the pathology of schizophrenia. In addition, cortical layers 2 and 4, and the cell-types must abundant in these layers, have become of particular interest to new research [5,10,11]. However, prior studies have produced inconsistent findings on the involvement of cortical layers and extent alterations throughout the brain.

Evidence from post-mortem studies consistently implicates impaired GABAergic interneurons in schizophrenia [4,6]. GABAergic interneurons are fundamental to neural circuits, exerting precise rhythmic inhibition on pyramidal cells and modulating neural oscillations [12,13]. Distinct interneuron subtypes can be identified based on their expression of calcium-binding proteins [13]—parvalbumin (PV), calbindin (CB), and calretinin (CR)—and neuropeptides, such as somatostatin (SST) [14]. Other cell-types, including vasoactive intestinal peptide (VIP), neuropeptide Y (NPY), and cholecystokinin (CCK), have been implicated in schizophrenia but research on these subtypes remains sparse [14,15].

While interneuron alterations can manifest at both the somal level and the presynaptic level, somal quantifications represent the most widely and systematically reported metrics across human post-mortem literature. Interneuron subtypes shape cortical information flow, are differentially expressed across cortical layers, and have been linked to schizophrenia pathophysiology [16–18]. Feedforward information enters layer 4 and propagates through layers 2/3 to higher-order areas [19], whilst feedback from higher-order areas returns via layer 1 and outputs from layers 5/6 [19]. *Bottom-up* accounts of schizophrenia propose a loss in precision of sensory inputs as key to the disorder. This manifests as deficits in sensory cortices [13] and feedforward-dominated layers (2/3/4) [19], with PV interneurons—the primary drivers of feedforward gamma-band oscillations [20]—as the key inhibitory substrate [21]. *Top-down* accounts emphasize failures in internal models originating in higher-order cortex [6,7] and feedback-dominated layers (1/5/6) [19], with proposed CR and SST interneuron involvement [22–26]. Altered *top-down* and *bottom-up* processing have both been proposed in schizophrenia [27], but more information at the level of the microcircuit is needed to clarify their involvement.

Current overviews of interneuron pathology in schizophrenia consist primarily of qualitative reviews [14,16] and meta-analyses limited to a single area or interneuron subtype [6]. A comprehensive meta-analysis across interneuron subtypes, cortical layers, and brain areas is therefore lacking. We addressed this gap by performing such an analysis in human post-mortem studies that examine interneuron deficits in schizophrenia and tests whether deficits are greatest in feedforward-dominated layers, in PV interneurons relative to *top-down*-associated subtypes, and whether alterations are modulated by the cortical hierarchy.

## METHODS

### Search strategy and selection criteria

We conducted a systematic review and meta-analysis following PRISMA guidelines [28]. Two investigators (A.G.M. and K.M.G.) independently screened all titles, abstracts, and full-text articles retrieved. Discrepancies at any stage were resolved via consensus or consultation with senior authors (P.J.U. and A.M.B.). ProQuest, PubMed, PsycINFO, Embase, and Google Scholar were searched from inception to Nov 24, 2025, with search terms relating to PV, CB, CR, and SST interneuron density and mRNA expression in schizophrenia (see Supplementary Methods for full search terms).

The primary outcomes for which data were sought were direct, quantitative, somal measures of GABAergic interneuron pathology, defined as: (1) interneuron cell density evaluated via immunohistochemistry (IHC) or immunocytochemistry, and (2) somatic mRNA expression across PV, SST, CB, and CR subtypes. Studies focusing exclusively on presynaptic bouton density, axonal arborizations, or non-subtype-specific enzymatic markers (e.g., GAD65/GAD67 puncta) were excluded. This boundary was established to maintain methodological homogeneity, as somatic counts and presynaptic terminal measures represent distinct cellular compartments with different scaling properties. Immunofluorescent labeling intensity or protein-level measures were not extracted because these index marker expression within cells instead of cell number and are not commensurable with IHC-based density measures. Where eligible studies reported both [17,29], we extracted only the somal density or mRNA measure. See Supplementary Materials for more information on the search strategy.

Studies were structurally grouped for statistical syntheses based on the neuroanatomical region evaluated, technical methodology (i.e., IHC-based cell density or mRNA expression levels), and interneuron subtype. Studies that used the same or overlapping samples were treated as independent when they analyzed different brain regions, cell-types, or methods (Table 1).

**Table 1.**
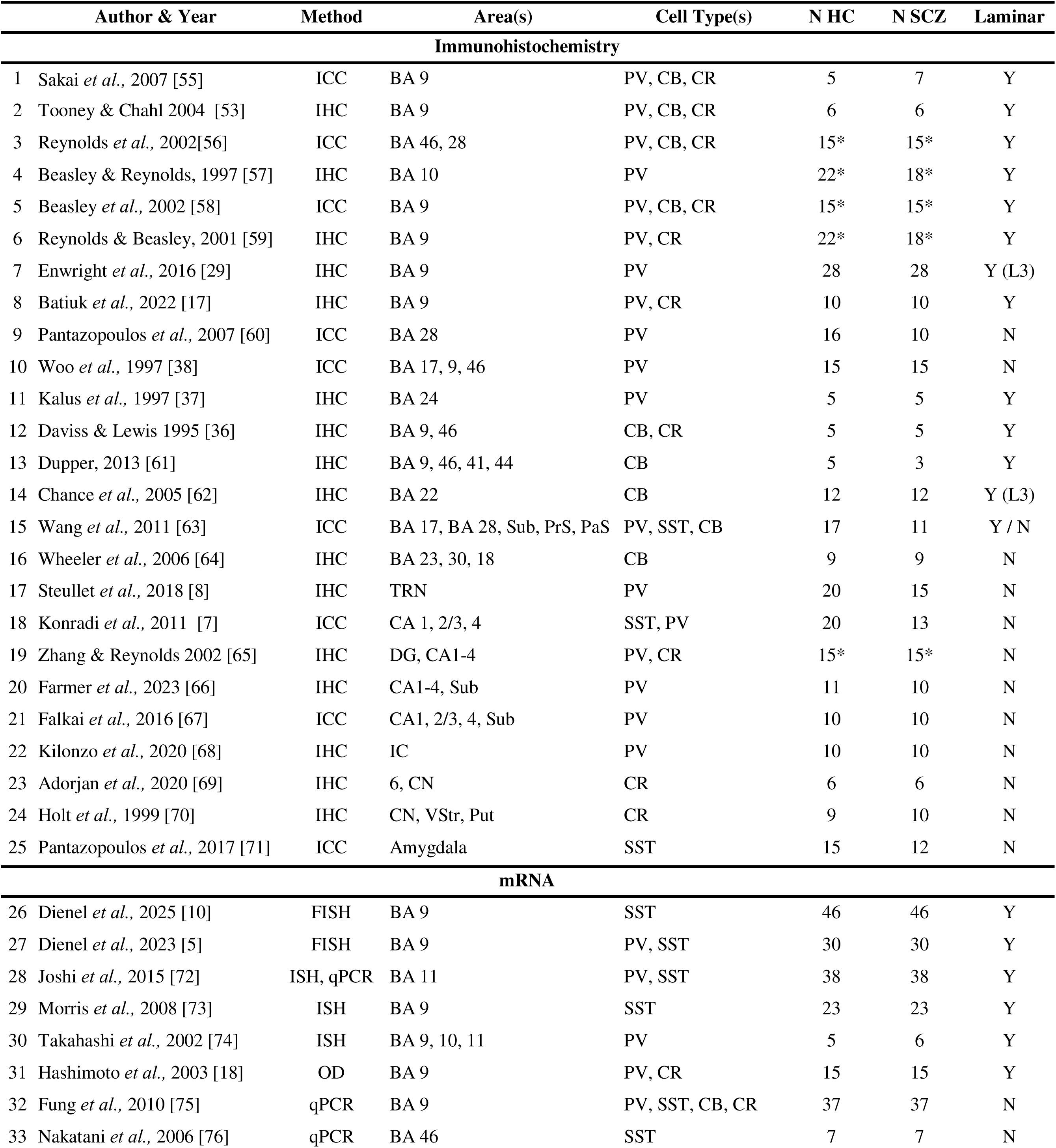
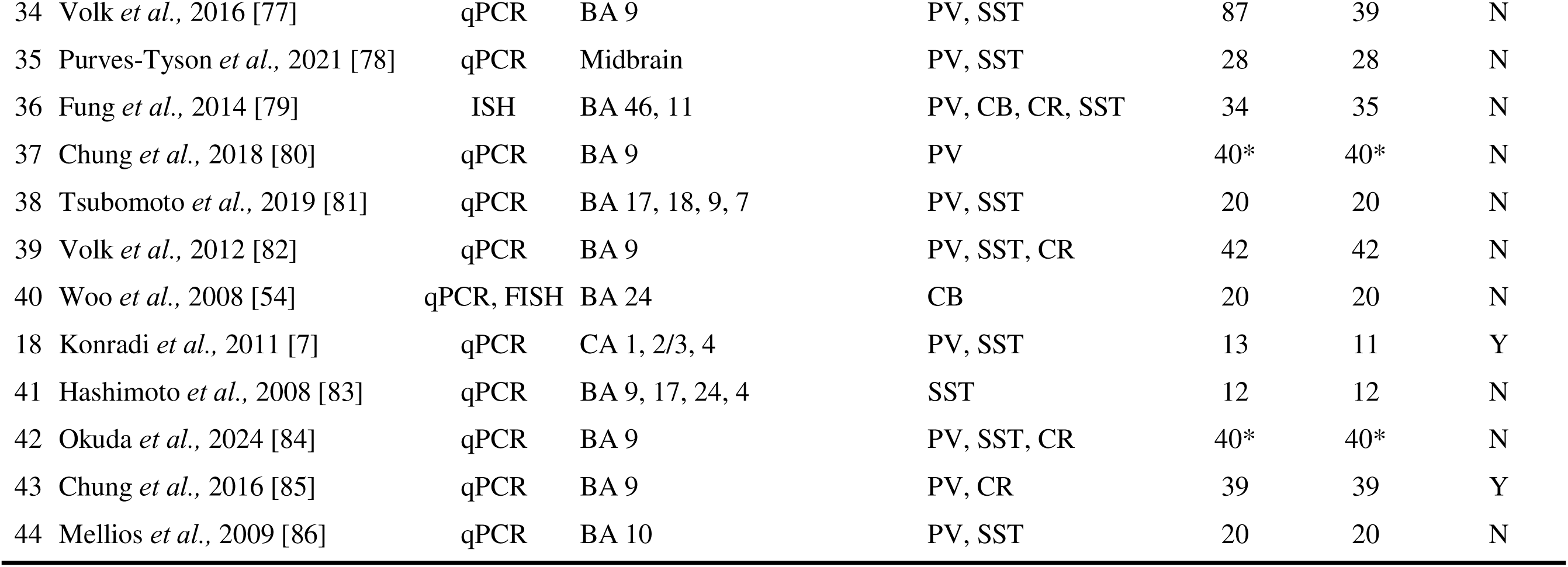
All papers included in the meta-analysis. First author, publication year, method(s), brain area(s), cell-type(s), number of subjects (N) for both healthy control (HC) and schizophrenia (SCZ) groups, if the study did (Y) or did not (N) examine cortical layers. (*) in the N columns indicate studies using the same or largely overlapping samples. BA=Brodmann area. ICC=immunocytochemistry. IHC=immunohistochemistry. FISH=fluorescent in situ hybridization. ISH=in situ hybridization. qPCR=quantitative polymerase chain reaction. OD=optical density. PV=parvalbumin. CB=calbindin. CR=calretinin. SST=somatostatin. Sub=subiculum. PrS=presubiculum. PaS=parasubiculum. TRN=thalamic reticular nucleus. IC=inferior colliculus. DG=dentate gyrus. CN=caudate nucleus. VSt=ventral striatum. Put=putamen. CA=Cornu Ammoni (hippocampal subfield). HC=healthy control. SCZ=schizophrenia.

Tables and Figure Captions
| Author & Year | Method | Area(s) | Cell Type(s) | N HC | N SCZ | Laminar |
| --- | --- | --- | --- | --- | --- | --- |
| **Immunohistochemistry** |  |  |  |  |  |  |
| 1 Sakai *et al.*, 2007 [55] | ICC | BA 9 | PV, CB, CR | 5 | 7 | Y |
| 2 Tooney & Chahl 2004 [53] | IHC | BA 9 | PV, CB, CR | 6 | 6 | Y |
| 3 Reynolds *et al.*, 2002[56] | ICC | BA 46, 28 | PV, CB, CR | 15* | 15* | Y |
| 4 Beasley & Reynolds, 1997 [57] | IHC | BA 10 | PV | 22* | 18* | Y |
| 5 Beasley *et al.*, 2002 [58] | ICC | BA 9 | PV, CB, CR | 15* | 15* | Y |
| 6 Reynolds & Beasley, 2001 [59] | IHC | BA 9 | PV, CR | 22* | 18* | Y |
| 7 Enwright *et al.*, 2016 [29] | IHC | BA 9 | PV | 28 | 28 | Y (L3) |
| 8 Batiuk *et al.*, 2022 [17] | IHC | BA 9 | PV, CR | 10 | 10 | Y |
| 9 Pantazopoulos *et al.*, 2007 [60] | ICC | BA 28 | PV | 16 | 10 | N |
| 10 Woo *et al.*, 1997 [38] | ICC | BA 17, 9, 46 | PV | 15 | 15 | N |
| 11 Kalus *et al.*, 1997 [37] | IHC | BA 24 | PV | 5 | 5 | Y |
| 12 Daviss & Lewis 1995 [36] | IHC | BA 9, 46 | CB, CR | 5 | 5 | Y |
| 13 Dupper, 2013 [61] | IHC | BA 9, 46, 41, 44 | CB | 5 | 3 | Y |
| 14 Chance *et al.*, 2005 [62] | IHC | BA 22 | CB | 12 | 12 | Y (L3) |
| 15 Wang *et al.*, 2011 [63] | ICC | BA 17, BA 28, Sub, PrS, PaS | PV, SST, CB | 17 | 11 | Y / N |
| 16 Wheeler *et al.*, 2006 [64] | IHC | BA 23, 30, 18 | CB | 9 | 9 | N |
| 17 Steullet *et al.*, 2018 [8] | IHC | TRN | PV | 20 | 15 | N |
| 18 Konradi *et al.*, 2011 [7] | ICC | CA 1, 2/3, 4 | SST, PV | 20 | 13 | N |
| 19 Zhang & Reynolds 2002 [65] | IHC | DG, CA1-4 | PV, CR | 15* | 15* | N |
| 20 Farmer *et al.*, 2023 [66] | IHC | CA1-4, Sub | PV | 11 | 10 | N |
| 21 Falkai *et al.*, 2016 [67] | ICC | CA1, 2/3, 4, Sub | PV | 10 | 10 | N |
| 22 Kilonzo *et al.*, 2020 [68] | IHC | IC | PV | 10 | 10 | N |
| 23 Adorjan *et al.*, 2020 [69] | IHC | 6, CN | CR | 6 | 6 | N |
| 24 Holt *et al.*, 1999 [70] | IHC | CN, VStr, Put | CR | 9 | 10 | N |
| 25 Pantazopoulos *et al.*, 2017 [71] | ICC | Amygdala | SST | 15 | 12 | N |
| **mRNA** |  |  |  |  |  |  |
| 26 Dienel *et al.*, 2025 [10] | FISH | BA 9 | SST | 46 | 46 | Y |
| 27 Dienel *et al.*, 2023 [5] | FISH | BA 9 | PV, SST | 30 | 30 | Y |
| 28 Joshi *et al.*, 2015 [72] | ISH, qPCR | BA 11 | PV, SST | 38 | 38 | Y |
| 29 Morris *et al.*, 2008 [73] | ISH | BA 9 | SST | 23 | 23 | Y |
| 30 Takahashi *et al.*, 2002 [74] | ISH | BA 9, 10, 11 | PV | 5 | 6 | Y |
| 31 Hashimoto *et al.*, 2003 [18] | OD | BA 9 | PV, CR | 15 | 15 | Y |
| 32 Fung *et al.*, 2010 [75] | qPCR | BA 9 | PV, SST, CB, CR | 37 | 37 | N |
| 33 Nakatani *et al.*, 2006 [76] | qPCR | BA 46 | SST | 7 | 7 | N |

**Table 1. All papers included in the meta-analysis.** First author, publication year, method(s),
|  |  |  |  |  |  |  |  |
| --- | --- | --- | --- | --- | --- | --- | --- |
| 34 | Volk *et al.*, 2016 [77] | qPCR | BA 9 | PV, SST | 87 | 39 | N |
| 35 | Purves-Tyson *et al.*, 2021 [78] | qPCR | Midbrain | PV, SST | 28 | 28 | N |
| 36 | Fung *et al.*, 2014 [79] | ISH | BA 46, 11 | PV, CB, CR, SST | 34 | 35 | N |
| 37 | Chung *et al.*, 2018 [80] | qPCR | BA 9 | PV | 40* | 40* | N |
| 38 | Tsubomoto *et al.*, 2019 [81] | qPCR | BA 17, 18, 9, 7 | PV, SST | 20 | 20 | N |
| 39 | Volk *et al.*, 2012 [82] | qPCR | BA 9 | PV, SST, CR | 42 | 42 | N |
| 40 | Woo *et al.*, 2008 [54] | qPCR, FISH | BA 24 | CB | 20 | 20 | N |
| 18 | Konradi *et al.*, 2011 [7] | qPCR | CA 1, 2/3, 4 | PV, SST | 13 | 11 | Y |
| 41 | Hashimoto *et al.*, 2008 [83] | qPCR | BA 9, 17, 24, 4 | SST | 12 | 12 | N |
| 42 | Okuda *et al.*, 2024 [84] | qPCR | BA 9 | PV, SST, CR | 40* | 40* | N |
| 43 | Chung *et al.*, 2016 [85] | qPCR | BA 9 | PV, CR | 39 | 39 | Y |
| 44 | Mellios *et al.*, 2009 [86] | qPCR | BA 10 | PV, SST | 20 | 20 | N |

Non-independence of samples was addressed via meta-regression. Importantly, recent evidence has established that no statistical differences between quantification techniques in mRNA (e.g., FISH or qPCR) exists [10], which we confirmed in our data set (see Supplementary Methods).

### Data extraction and synthesis

For data represented in plots, means and variances were extracted using WebPlotDigitizer [30]; otherwise, numerical data were extracted directly from manuscripts. For each study, we extracted the mean (*x*-), variance, and sample size (*n*) for both the control and schizophrenia groups across all reported cell types, brain areas, and cortical layers. Missing covariate details were managed via pairwise deletion in meta-regressions. All statistical analyses and data visualization were performed in MATLAB [31], except for the visualization of brain plots (see Supplementary Methods). We adopted a random-effects analytical model, using sample size, population mean, and standard deviation (SD) to calculate Hedges’ *g* effect size (equations 1, 2) and sampling variance of *g* (equation 3) for each observation.

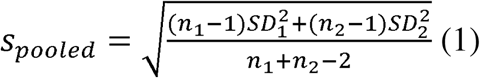

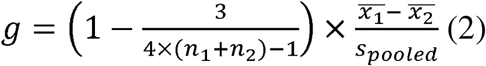

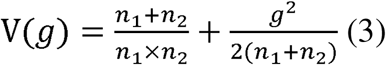

We aggregated data across several datasets (i.e., PV/CB/CR/SST + IHC/mRNA) and reported mean *g* with the *p*-value from a one-sample t-test to assess significance of datasets. Where appropriate, *p-*values were corrected for multiple comparisons using Benjamini-Hochberg False Discover Rate (FDR) [32], indicated by *p_FDR_* (see Supplementary Methods, Supplementary Table 6). To compare the extent of effects across areas, interneurons, and datasets, we used a two-sample t-test and Cohen’s effect sizes analysis. Comparisons were also FDR-corrected (Supplementary Table 6). When available, Hedges’ *g* was computed for each cortical layer; if a study combined layers (e.g., layers 5/6), the effect size was used for both layers (Supplementary Methods). For studies without laminar specificity, a single effect size was calculated from the mean and SD of each area and/or cell-type. The number of observations (*n_o_*) refers to multiple datapoints for the same group. For example, primary and secondary visual cortices (V1 and V2, respectively) from one study may be analyzed as unique datapoints within the visual cortex (VCx). Otherwise, the number of studies (*n_s_*) refers to combining areas in the same general structure (e.g., V1/V2 in VCx). This allows us to examine alterations in unique brain areas arising from the same study (*n_o_*) and also perform sensitivity analyses across studies (*n_s_*). Where appropriate, we report both *n_o_* and *n_s_*.

### Linear mixed-effects modelling

Prior to synthesis, data eligibility was determined based on the completeness of reporting; nested comparisons were extracted at the finest anatomical resolution provided (by specific layer or region). In instances where studies reported data split across multiple fields or cortical layers without a consolidated cohort mean, raw values were pooled using standard sample-weighted variance combining formulas to ensure structured data alignment (Supplementary Methods). To synthesize the effect sizes whilst accounting for heterogeneity across studies, risk of bias, brain regions, cell-types, methods, sample sizes, and cortical layers, we employed linear mixed-effects models (LMMs). LMMs were chosen because they explicitly model hierarchical data structure (e.g., multiple observations per brain area) and non-independence (e.g., repeated measures from the same tissue bank), while allowing flexible testing of fixed-effects and targeted contrasts for our hypotheses.

Categorical variables were parameterized with effects coding. We employed effects coding instead of the default reference coding to remove arbitrary selection references. Consequently, the model’s intercept (*β*_0_) represents the grand mean of the effect size averaged across all levels, *K*, of the categorical variables (e.g., cell-type, brain area, and layer) and adjusted for other covariates. Thus, each fixed-effect estimate (*β_nk_*) represents that variables’ deviation from the grand mean (*Σ*), *g_ij_* represents the observed effect size for observation *j* within cluster *i* (repository/study), *β*_0_ represents the fixed-effect intercept, b_0*i*_ represents the random intercept for cluster *i*, *β_n_* represents the fixed-effect coefficient for the predictors, and *∈_ij_* represents the residual error.

We extracted demographic data, where available, from each study (Supplementary Table 1). We performed a meta-regression (Model 1) on subject age, sex (assigned at birth), post-mortem interval (PMI), brain pH, sample size (*N*), cell-type, method (IHC or mRNA), and area grouping (PFC, hippocampus, non-PFC cortex, or subcortex) (see Supplementary Methods for details regarding age, sex, PMI, and pH calculations). We also modeled fixation method and, because individual subject IDs were not consistent or identifiable across studies, we included study and tissue repository as random intercepts to account for any potential inter-study or inter-repository variability. We report the *F*-statistic and FDR-corrected *p*-value from an ANOVA, as well as the post-hoc fixed-effects coefficient (β), *t*-statistic, standard error (SE), and *p*-value from the model.

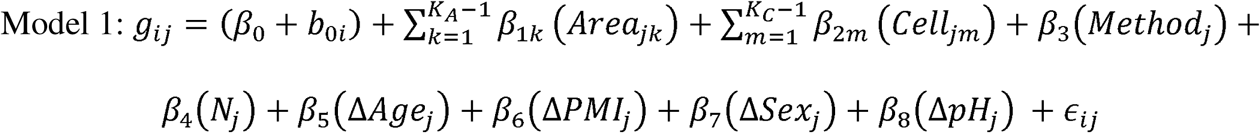

To derive denoised effect sizes, we employed two additional LMMs with effects coding for categorical fixed-effects. Both models included cell-type and brain area as fixed-effects and methodology and sample size as fixed covariates (Model 2). Model 3 included cortical layer as an additional fixed-effect. These models allowed us to examine deficits in target variables (e.g., cell-type, area, or layer), while controlling for nuisance variables (e.g., method and sample size) and non-independence (e.g., study and repository). Methodology (i.e., IHC or mRNA) was included in the model as a categorical covariate to control for differences between methods and focus results on cellular-level alterations. We performed exploratory analyses on IHC versus mRNA effect sizes for each cell-type, cell-type specific changes across brain areas, cell-type specific changes across cortical layers (Supplementary Analyses).

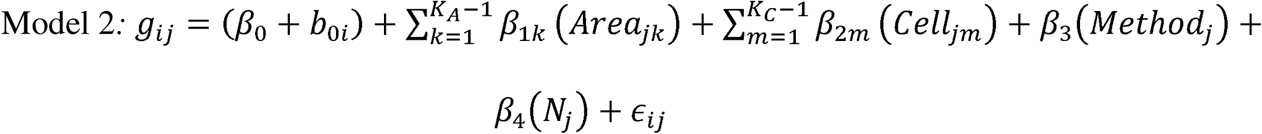

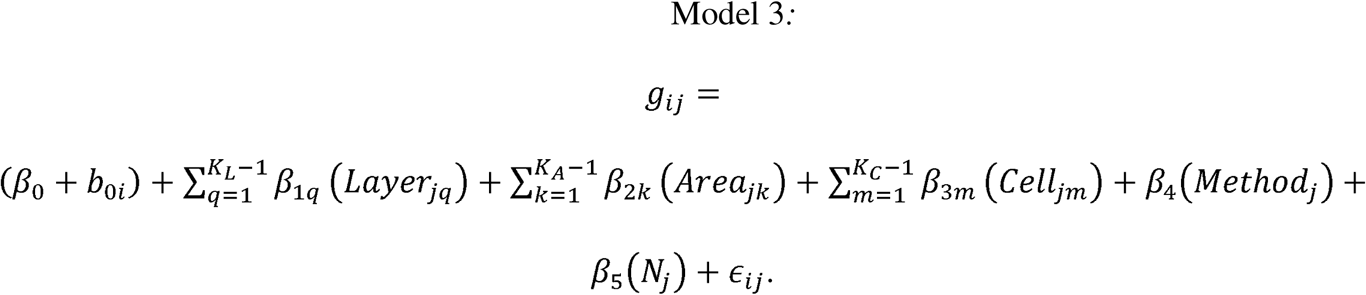

We used the fixed-effects coefficients from the model to calculate the estimated marginal means (EMMs) [33], which represent Hedges’ *g* after adjusting for other covariates and clustering by the random-effects. The EMM of PV, for example, is calculated according to equation 4. Significance derived from the model (*p_β_*) indicates a significant deviation from the global mean measured by that model. We also assessed significance of the EMM from zero (*p_0_*) with Wald *F*-tests to determine if the target variable exhibited deficits in schizophrenia compared to controls. These values were FDR-corrected (*p*_0,*FDR*_).

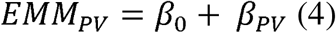

### Hypothesis testing

We perform planned linear contrasts on the EMMs to address three questions regarding the *top-down* versus *bottom-up* hypothesis: (1) are *top-down* or *bottom-up* layers more affected; (2) are *top-down* or *bottom-up* cell-types more affected; (3) do effect sizes scale with the cortical hierarchy? For cortical layers, we compared all *bottom-up* layers (2/3/4) versus all *top-down* layers (1/5/6), *bottom-up* input layer 4 versus *top-down* input layer 1, and *bottom-*up output layers 2/3 versus *top-down* output layers 5/6. For cell-types, we compared PV against CR, SST, CB/SST combined, and CR/CB/SST combined. For brain areas, we tested whether EMMs scaled with the hierarchy defined by Barrett and Miller [34], which organizes included areas into posterior areas (visual cortex [VCx], primary auditory cortex [A1], posterior parietal cortex [PPC]), anterior areas (PFC, anterior and posterior cingulate cortices [ACC, PCC], Broca’s area [BrA], primary motor cortex [MCx]), entorhinal cortex (EC), and hippocampus groups. We report the difference in EMMs (dEMM) between target groups. Significance of these contrasts was determined using Wald *F*-tests and subsequently corrected with FDR. The difference between PV and other cell-types, for example, would be computed according to equation 5.

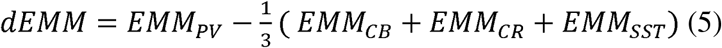

### Sensitivity analyses

Publication bias was explored quantitatively using Egger’s linear regression tests, with correction for multiple comparisons [35]. Funnel plots were constructed for each dataset to visually inspect bias. We assessed heterogeneity by calculating *I*^2^ and *τ*^2^ for each dataset for area grouping (i.e., PFC, hippocampus, non-PFC cortex, subcortex) where *n_s_*≥5. We also explored whether this heterogeneity could be explained by differences in number of studies across the sample using LMM Model 1.

### Risk of bias assessment

Risk of bias for each study was evaluated by two reviewers (A.G.M. and K.M.G.), tracking demographic and tissue quality matching between schizophrenia and healthy control cohorts. Discrepancies were resolved through consensus. The overall certainty and robustness of the synthesized evidence were systematically evaluated using ‘leave-one-author-out’ sensitivity validation, iteratively removing single study cohorts to confirm that omnibus LMM parameter estimates were stable and not influenced by a particularly large sample size or effect size. We implemented a *t*-test across leave-one-out iterations to determine stability.

### Code Availability

Codes for the analyses were written in MATLAB R2024a (The MathWorks Inc., Natick, MA) and can be found at https://github.com/BastosLab/sim-map, along with pre-processed data.

## RESULTS

### Sample

Out of the total 1318 records screened, 44 studies were included (Fig. 1A). The final sample consisted of 814 healthy controls (HCs) and 736 individuals with schizophrenia (Table 1, Supplementary Table 1). PFC and hippocampus were the predominantly quantified areas, appearing in 73% of all studies (Fig. 2, Supplementary Table 3). PV was the most studied cell-type (*n_s_=*32 studies), followed by SST (*n_s_* =17), CR (*n_s_*=16), and CB (*n_s_*=13).

**Fig. 1.**
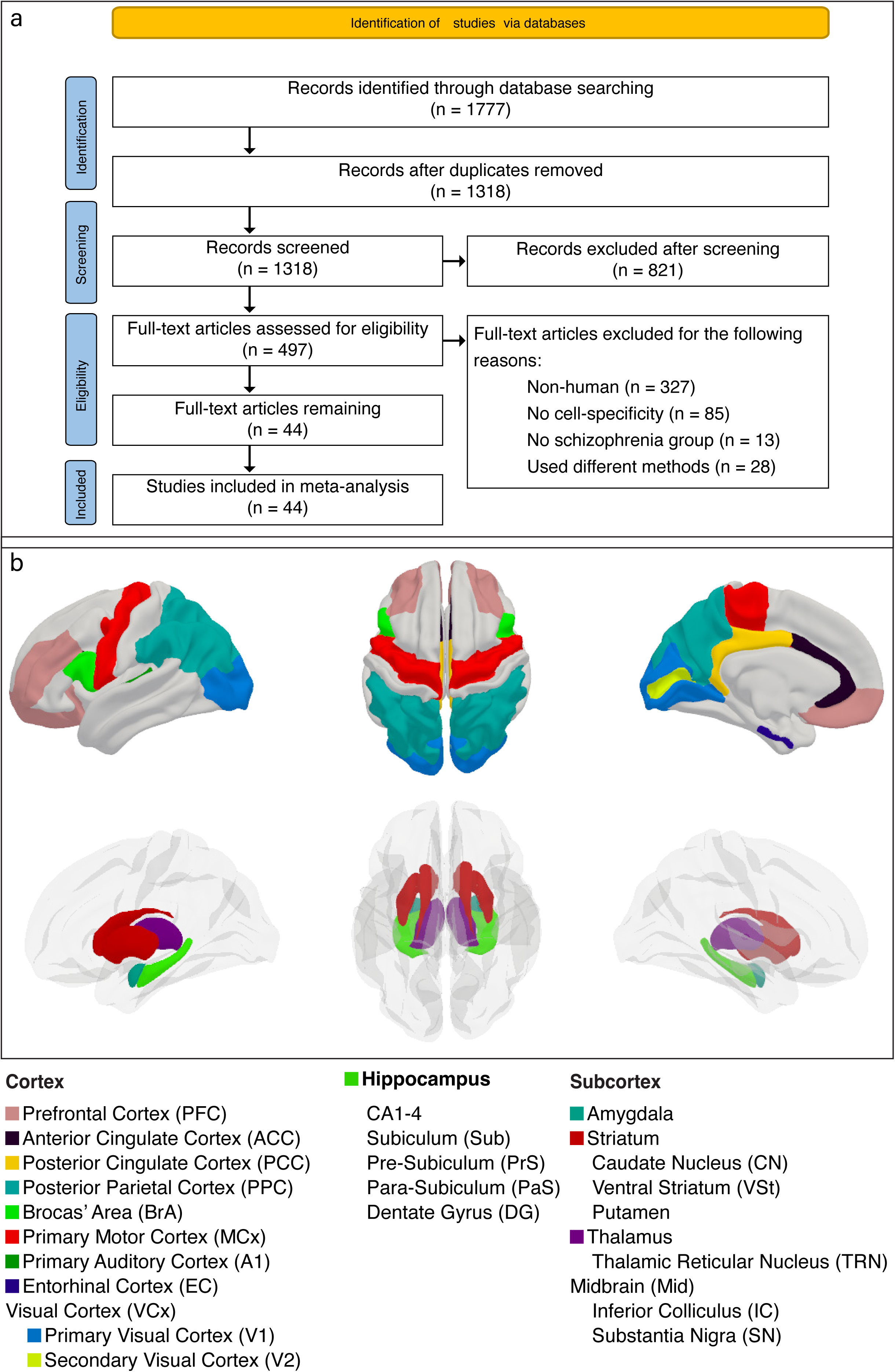
PRISMA flowchart and brain areas sampled. (**a**) PRISMA 2020 flowchart for study screening and inclusion/exclusion. (**b**) Each included brain region is shaded in its own designated color. Hippocampal, striatal, and thalamic subregions are not shown. Also not depicted are the midbrain regions, which are not included in the utilized atlas. See Supplementary Methods for information regarding mapping our data onto brain plots.

**Fig. 2:**
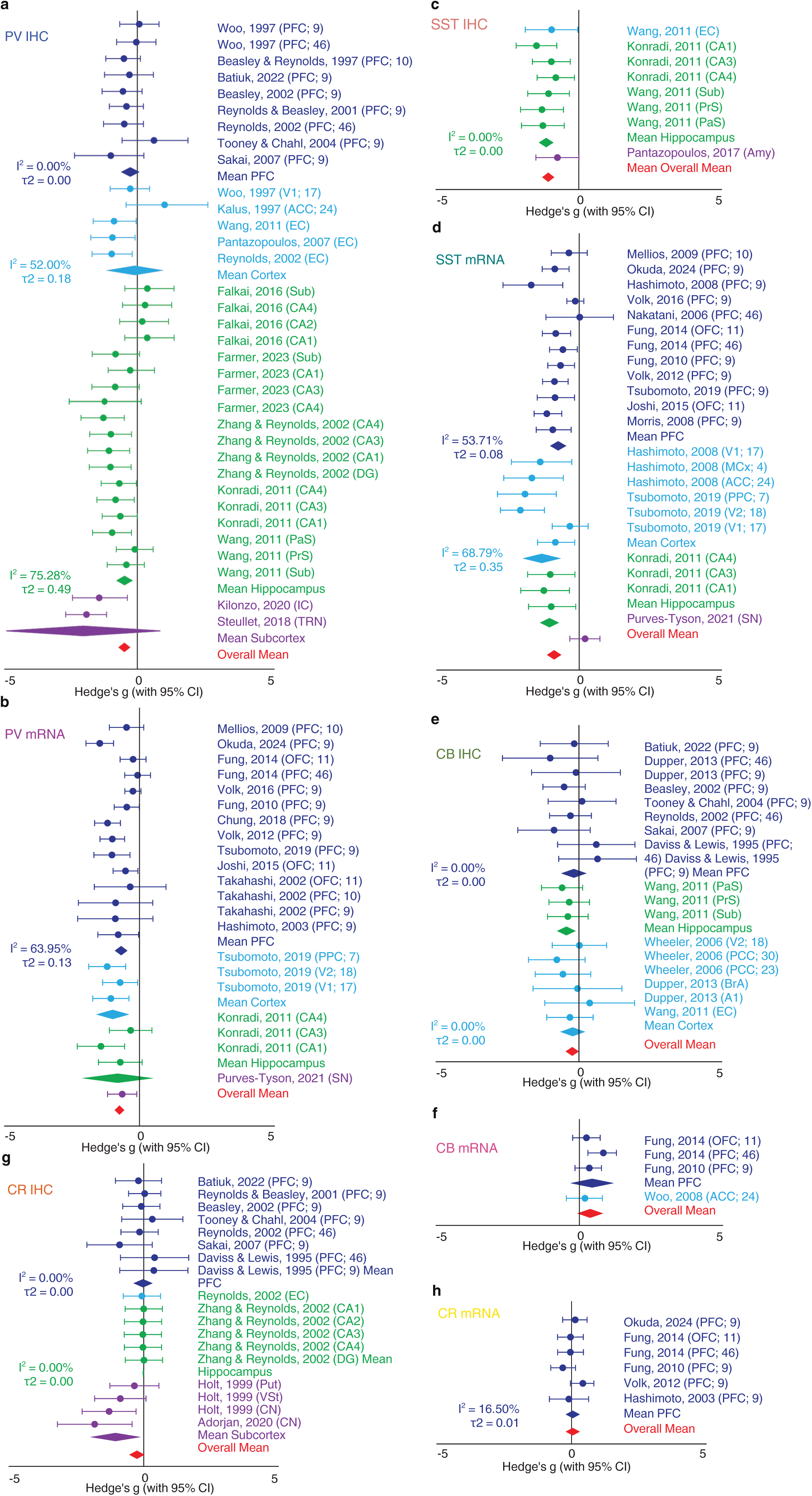
Forest plot of Hedges’ g effect sizes across studies. (**a**) PV IHC and (**b**) PV mRNA studies; (**c**) SST IHC and (**d**) SST mRNA studies; (**e**) CB IHC and (**f**) CB mRNA studies; and (**g**) CR IHC and (**h**) CR mRNA studies. Error bars represent 95% confidence intervals (CIs). Where data were sufficient, group means are also shown for PFC, non-PFC cortical areas, hippocampus, subcortex, and the overall mean across that cell-type/method group. I^2^ and *τ*^2^ are also given when there were sufficient studies to perform this calculation (n_s_≥4). To improve visualization, one outlier (CA2 from Zhang & Reynolds, 2002) was removed from subpanel (a) PV IHC (*g*=–6.011). Note, this study was not biased nor was it excluded from data analyses.

### Regional alterations

We first considered IHC studies across layers and brain subregions, which revealed deficits of IHC-based interneuron density (density) in hippocampus for PV (*g*=–0.81, *n_o_*=19, *n_s_*=5, *p*=0.020, Fig. 3G) with high heterogeneity (*I*^2^=75.28%, Fig. 2A, Supplementary Table 4). There were also significant reductions in density in hippocampus for CB (*g*=–0.44, *n_o_*=3, *n_s_*=1, *p*=0.029, Fig. 2E, 3J), and SST (*I*^2^=0.00%, Fig. 2C; *g*=–1.16, *n_o_*=6, *n_s_*=2, *p*<0.0001, Fig. 3H), as well as PV deficits in EC (*I*^2^=52.00%, Fig. 2A; *g*=–0.54, *n_o_*=3, *n_s_*=3, *p*=0.0026, Fig. 3G) and CR deficits in the striatum (*g*=–1.10, *n_o_*=4, *n_s_*=2, *p*=0.038, Fig. 2G, 3I). PV deficits in EC (*p*_FDR_=0.012) and SST in hippocampus (*p*_FDR_<0.001) remained significant after FDR-correction.

**Fig. 3:**
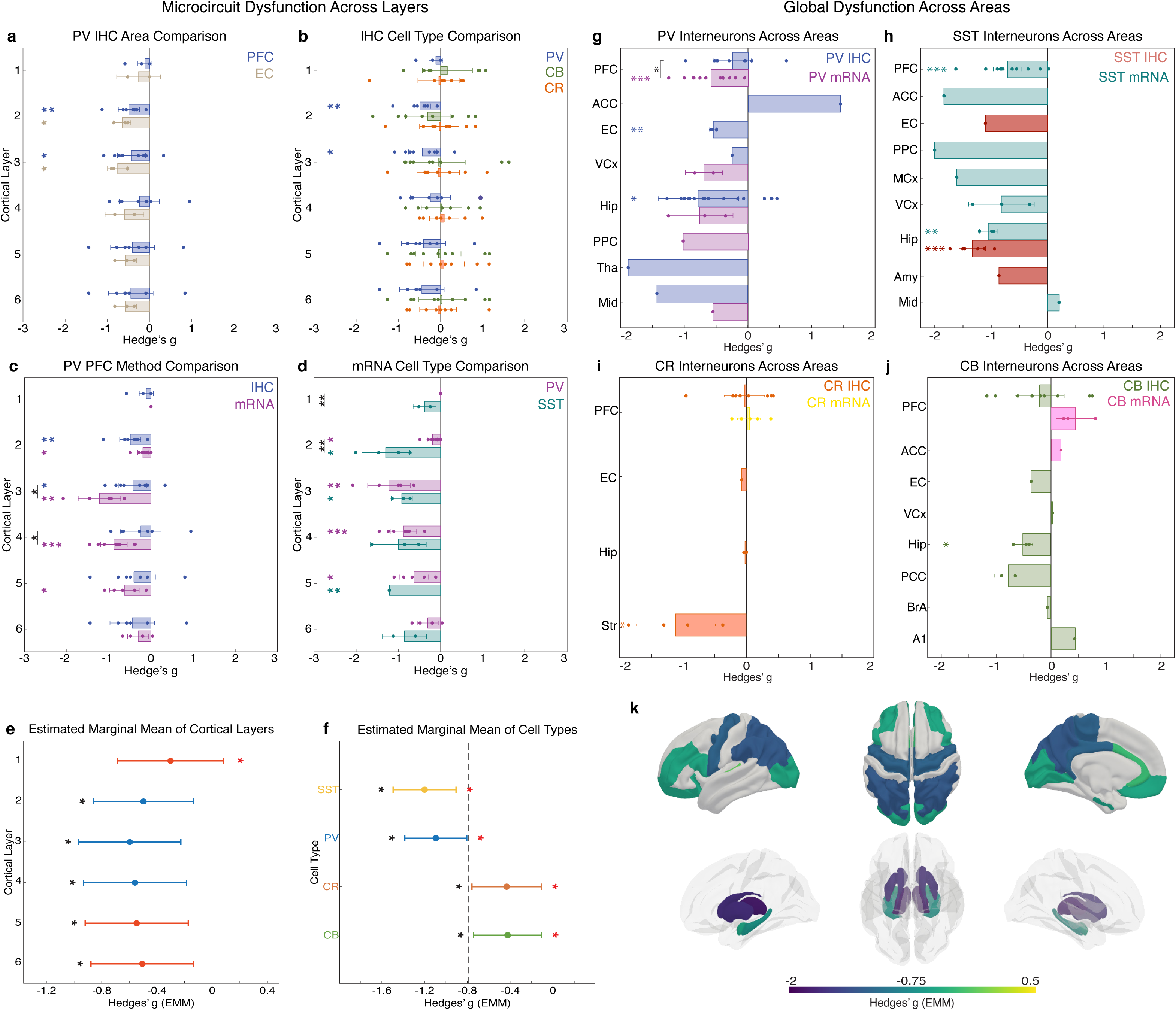
Microcircuit dysfunction across cortical layers and global dysfunction across brain areas. (**a-d**) Laminar bar plots with specific comparisons for IHC (cell density) (**a**) PFC and Entorhinal Cortex (EC) and (**b**) PV, CB, and CR interneurons in the PFC. Bar plots for mRNA expression comparing (**c**) IHC and mRNA for PV interneurons in the PFC and (**d**) PV and SST interneuron expression in the PFC. Colored dots are individual observations, with the mean +/− 2 standard errors of the mean. Significance of each bar indicated with color asterisks. Comparisons within each layer are shown as black asterisks with black lines. \*\*\**p*<0.001, \*\**p*<0.01, \**p*<0.05. (**e**) Estimated marginal means (EMMs) for cortical layers across cell-types and brain areas and (**f**) cell types across brain areas from Model 2. Error bars represent 95% CI, with a black asterisk representing a significant deviation from 0, and a red asterisk indicating a significant deviation from the global mean (dashed line). (**g-j**) Bar plots for each area and method for (**g**) PV, (**h**) SST, (**i**) CR, and (**j**) CB interneurons. If an area has data from both IHC and mRNA, both are shown in distinct colors. Significance of each bar via one-sample t-test is indicated with color asterisks. Significant difference between groups where *n*>1 is computed with a paired t-test and shown as black bracket with black asterisks. \*\*\**p*<0.001, \*\**p*<0.01, \**p*<0.05. (**k**) Heatmaps of each brain area’s EMM with color corresponding to the size of the effect. Midbrain is not depicted (EMM=–0.171). Areas are shown equally in both hemispheres, though laterality was not explored.

We also considered mRNA studies across layers and brain subregions, which revealed significant reductions of PV in PFC (*I*^2^=63.95%, Fig. 2B; *g*=–0.68, *n_o_*=14, *n_s_*=11, *p*<0.0001, Fig. 3G), and SST PFC (*I*^2^=53.71%, Fig. 2D; *g*=–0.71, *n_o_*=12, *n_s_*=11, *p*<0.001, Fig. 3H) and in hippocampus (*I*^2^=68.79%, Fig. 2D; *g*=–1.04, *n_o_*=3, *n_s_*=1, *p*=0.0056, Fig. 3H), all of which remained significant after FDR-correction (*p*_FDR_<0.05).

Within PFC, PV mRNA expression was more reduced than CB (Cohen’s *d*=–2.73, 95% CI: [–4.23, –1.17], *p*<0.001) and more reduced than CR (Cohen’s *d*=–1.89, 95% CI: [–2.97, – 0.77], *p*<0.001), and SST was more reduced than CR (Cohen’s *d*=–1.85, 95% CI: [–2.96, –0.70], *p*=0.0013). Within IHC studies of the hippocampus, CB was more reduced than CR (Cohen’s *d*=–4.71, 95% CI: [–7.55, –1.81], *p*<0.001), SST was more reduced than CR (Cohen’s *d*=–5.53, 95% CI: [–8.24, –2.78], *p*<0.0001), and SST was more reduced than CB (Cohen’s *d*=–2.84, 95% CI: [0.90, 4.71], *p*=0.0027). All cell-cell comparisons remained significant after FDR-correction (all *p*_FDR_<0.01). SST mRNA expression deficits were greater in non-prefrontal cortical areas than in PFC (Cohen’s *d* = 1.13, 95% CI: [0.10, 2.12], *p* = 0.031), and PV mRNA expression was significantly more reduced than density in PFC (Cohen’s *d* = 0.98, 95% CI: [0.11, 1.83], *p* = 0.027); however, these comparisons were not significant after FDR-correction (*p*_FDR_>0.05).

The majority of studies (82%) maintained a low-to-moderate risk of bias due to rigorous case-control matching for primary demographics (age and sex) and PMI, and other factors (Supplementary Table 1, 2). A higher risk of bias was predominantly confined to historical cohorts [36–38] where reporting on technical and demographic factors was incomplete or sample sizes were small. Egger’s regression did not reveal any publication bias after FDR correction, although some cell-type datasets were visually asymmetric (Supplementary Fig. 1). Importantly, omnibus ANOVA coefficients were stable across leave-one-author-out iterations (Supplementary Table 5). Aside from PV IHC in hippocampus, heterogeneity remained low-to-moderate for all other datasets tested (Supplementary Table 4).

### Cortical layer alterations

We also explored effects at the level of cortical layer. PV interneuron density in PFC and entorhinal cortex (EC) layer 2 (PFC: *g*=–0.49, *n_o_*=7, *n_s_*=7, *p*=0.0079; EC: *g*=–0.65, *n_o_*=3, *n_s_*=3, *p*=0.025) and layer 3 (PFC: *g*=–0.43, *n_o_*=8, *n_s_*=8, *p*=0.036; EC: *g*=–0.75, *n_o_*=3, *n_s_*=3, *p*=0.025) was reduced in schizophrenia compared to controls (Fig. 3A). CB and CR interneuron densities were not affected in any PFC layer (all *p*>0.05, Fig. 3B), and insufficient data precluded laminar analysis in other areas.

For PFC mRNA expression, PV was reduced in layers 2 (*g*=–0.20, *n_o_*=7, *n_s_*=5, *p*=0.016), 3 (*g*=–1.22, *n_o_*=5, *n_s_*=3, *p*=0.0083), 4 (*g*=–0.92, *n_o_*=7, *n_s_*=5, *p*<0.001), 5 (*g*=–0.63, *n_o_*=5, *n_s_*=3, *p*=0.023) (Fig. 3C). SST mRNA expression was reduced in layers 2 (*g*=–1.25, *n_o_*=4, *p*=0.030), 3 (*g*=–0.85, *n_o_*=3, *p*=0.044), and 5 (*g*=–1.21, *n_o_*=2, *p*=0.0016; Fig. 3D). After FDR correction, only PV mRNA in PFC layer 4 (*p*_FDR_=0.019) and SST mRNA in PFC layer 5 (*p*_FDR_=0.029) were significantly reduced.

PV mRNA studies exhibited significantly larger effect sizes than PV IHC studies in layers 3 (Cohen’s *d*=1.46, 95% CI: [0.23, 2.63], *p*=0.019) and 4 (Cohen’s *d*=1.24, 95% CI: [0.12, 2.32], *p*=0.029; Fig. 3C). SST mRNA expression was more affected than PV mRNA in layer 1 (Cohen’s *d*=3.69, 95% CI: [1.044, 6.26], *p*=0.0033; Fig. 3D) and 2 (Cohen’s *d*=2.44, 95% CI: [0.83, 3.99], *p*=0.0021; Fig. 3D). None of the Cohen’s comparisons remained significant after FDR correction (*p*_FDR_>0.05).

### Linear mixed-effects modelling

The above results are based on pooled and aggregated Hedge’s g values across studies and observations in the literature. In addition, we systematically used Linear mixed-effects modeling (LMMs) to model the effects of method, demographic factors, including number of subjects, differences in age, PMI, as well as technical factors such as tissue repository, fixation method, reagent manufacturer, and other factors (see Methods and Supplemental Tables 1, 7) in mediating the observed effects. Here we report the the Estimated Marginal Means (EMMs) of the LMMs, which are effect size estimates that account for all modeled covariates and therefore separate the main effects of interest (e.g., PV or SST effect size) from these other nuisance factors.

LMMs revealed significant main effects of cell-type (Models 2 and 3, both *p_FDR_*<0.0001), brain area (Model 2 *p_FDR_*=0.041; Model 3 *p_FDR_*=0.022), and sample size (Model 2 *p_FDR_*=0.0029; Model 3 *p_FDR_*=0.0010). All four cell-types were significantly reduced compared to zero (Model 2, grand mean EMM=−0.79; Fig. 3F). SST interneurons (EMM=–1.20, *p_0,FDR_*<0.0001) were the most reduced, followed by PV (EMM=–1.10, *p_0,FDR_*<0.0001). CB (EMM=–0.42, *p_0,FDR_*=0.010) and CR (EMM=–0.43, *p_0,FDR_*=0.010) were significantly reduced compared to zero but less so than PV and SST. Note that some of these effects, particularly for CB and CR, were different from the reported effects based on simple aggregates of Hedge’s g values (e.g., shown in Figure 2). The LMM models had in some cases different effects because they controlled for the effect of area, *N*, and methodology, suggesting they provided stronger power in the analysis and could reveal previously hidden effects.

Examining brain areas, the thalamus showed the largest deficit (EMM=–1.80, *p_0,FDR_*<0.001) and primary auditory cortex the least (EMM=–0.097, *p_0,FDR_*=0.90). PFC (EMM=–0.52, *p_0,FDR_*<0.0001) and hippocampus (EMM=–0.69, *p_0,FDR_*<0.001) were both significantly reduced but were not among the most affected areas (Fig. 3K). Each cell-type also exhibited unique patterns of alterations across areas that generally followed the same trends here (Supplementary Fig. 4).

Cortical layer was also a significant predictor of effect size (*p_FDR_*=0.049, Fig. 3E). Post-hoc analyses revealed that only layer 1 did not significantly deviate from zero (EMM=–0.32, *p_0,FDR_*=0.13, Fig. 3E). Layer 3 was the most affected layer (EMM=–0.62), followed by L4 (EMM=–0.58), L5 (EMM=–0.56), L2 (EMM=–0.54), and L6 (EMM=–0.52) (all *p_0,FDR_*<0.01). In an interaction model, SST interneurons were significantly reduced in L1-6, while PV interneurons were reduced in L4. CB and CR interneurons were not affected in any layer (note that the data for this analysis were limited to only area PFC), which aligns with our findings on the laminar resolution of these interneuron subtypes (Supplementary Fig. 2, 5).

### Hypothesis testing

Planned linear contrasts revealed that *bottom-up* input layer 4 was significantly more affected than *top-down* input layer 1 (dEMM=–0.26, 95% CI: [–0.45, –0.079], *p_FDR_*=0.017; Fig. 4C). The combined *bottom-up* layers (layers 2/3/4) were more affected than the *top-down* layers (layers 1/5/6, dEMM=–0.11, 95% CI: [–0.21, –0.012], *p_FDR_*=0.044), and no significant difference was found between output layers (L2/3 vs L5/6). PV interneurons were significantly more affected than CR (dEMM=–0.66, 95% CI: [–0.91, –0.42], *p_FDR_*<0.0001; Fig. 4D), CB/SST combined (dEMM=–0.29, 95% CI: [–0.48, –0.093], *p_FDR_*=0.016; Fig. 4D), and all non-PV types combined (dEMM=–0.41, 95% CI: [–0.59, –0.24], *p_FDR_*<0.001; Fig. 4D). PV and SST interneurons were not statistically different (dEMM=0.099, 95% CI: [–0.11, 0.31], *p_FDR_*=0.78). Brain area was a significant predictor of effect size (*p_FDR_*=0.041), but deficits did not scale with the cortical hierarchy (Fig. 3K, 4E; Supplementary Fig. 3, 4). All hierarchical groups exhibited significant reductions: anterior areas (EMM=–0.74, 95% CI: [–1.21, –0.28], *p_0,FDR_*=0.0046), hippocampus (EMM=–0.69, 95% CI: [–0.98, –0.40], *p_0,FDR_*<0.0001), posterior areas (EMM=–0.63, 95% CI: [–1.20, –0.060], *p_0,FDR_*=0.0034), and EC (EMM=–0.55, 95% CI [–0.96, –0.14], *p_0,FDR_*=0.012).

**Fig. 4:**
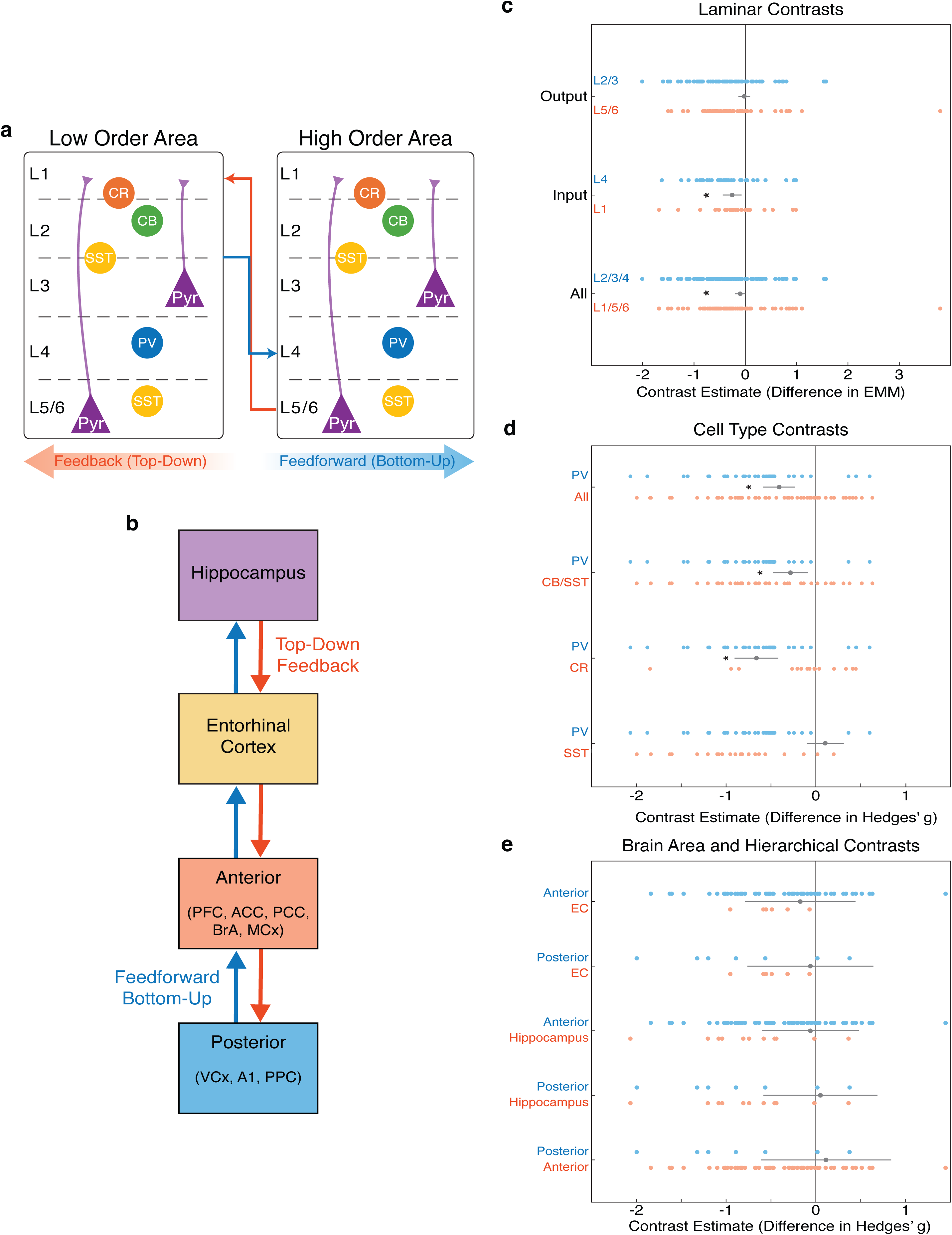
Hypothesis testing. Estimated marginal means (EMMs) from linear mixed effects models, as well as the planned linear contrasts for hypothesis testing. **(a)** Schematic of top-down and bottom-up communication between cortical layers and interneuron cell-types in low-order and high-order areas. Red arrows indicate feedback (*top-down*) communication from layers L5/6 to L1. Blue arrows indicate feedforward (*bottom-up*) communication from layers L2/3 to L4. Pyramidal cells are labelled Pyr. (**b**) Hierarchical schematic of cortical areas. (**c–e**) Planned linear contrasts for cortical layers (**c**), cell-types (**d**), and brain areas/hierarchical levels (**e**). In all contrast panels, the bottom-up side of each contrast is shown in blue dots and the top-down side in red dots. The difference between groups (contrast estimate) is shown as a grey dot with 95% confidence interval (CI). Significance is indicated by black asterisks when the contrast estimate differed significantly from 0, as assessed with a Wald F-test (*p*<0.05).

### Sensitivity analyses

To ensure that our findings were not artifactually driven by demographic or technical factors, we used a meta-regression (Model 1) that controlled for such variables. An omnibus ANOVA revealed that age (*p_FDR_*=0.46), sex (*p_FDR_*=0.74), PMI (*p_FDR_*=0.46), pH (*p_FDR_*=0.50), methodology (*p_FDR_*=0.46), and area (*p_FDR_*=0.46) were not significant drivers of effect size. We did identify a significant effect of sample size (*p_FDR_*=0.033). Despite being a very small effect (β=0.0039), we embedded sample size as a fixed covariate to regress out this effect. In Model 1, we also identified a robust overall (averaging across area/layer) effect of cell-type (*F*(3,42)=14.23, *p_FDR_*<0.0001), which confirms that alterations in cell-types is robust to the inclusion of substantial technical factors. The random intercepts of study, tissue repository, and fixation method were negligible (*σ*^2^<0.0001). There was insufficient data to include the effect of medication (Supplementary Table 1). In a separate analysis, we confirmed that reagent manufacturer did not significantly modulate effect size (Supplementary Analyses, Supplementary Table 7).

## DISCUSSION

We provided an overview of the specific interneurons, cortical layers, and brain areas implicated in aberrant GABAergic neurotransmission in schizophrenia. We performed a comprehensive meta-analysis of post-mortem interneuron data, which included studies quantifying both interneuron density and mRNA expression levels. We found that PV and SST interneuron density and expression were the most consistently reduced, whilst CB and CR interneurons were less affected. In addition, layers 3 and 4 exhibited the largest deficits, regardless of cell-type, which aligns with prior work [17], though areas with laminar resolution were limited to PFC and EC. Future research should consider analyzing deficits at the level of cortical layer across the brain. Among the brain areas implicated, PFC and hippocampus were consistently affected but less so than subcortical structures, such as the thalamus and striatum.

These cellular and laminar alterations reflect disruptions embedded in the fundamental architecture of the cortical column that could explain neurophysiological signatures of schizophrenia that in turn have relevance for symptom generation and treatment development. PV and SST interneurons work cooperatively to produce rhythmic gamma oscillations [39], linked to altered information processing in schizophrenia [21]. Moreover, optogenetic stimulation of PV interneurons improves schizophrenia-like cognitive rigidity in animal models [40], likely through recovery of PV-generated gamma. Selective and targeted modulation of these interneurons is, therefore, an underexplored but potentially fruitful avenue for treatment development and drug discovery [41]. The clinical relevance of GABAergic interneuron deficits is underscored by recent evidence that reduced *PVALB* expression—the gene that encodes PV interneurons—can be detected via peripheral blood samples in schizophrenia [42]. By identifying PV and SST interneurons in layers 3/4 as the principal sites of pathology, our work highlights these cells as high-priority targets for pharmacological rescue with GABAergic modulators, whilst also providing a clear mechanistic foundation for non-invasive gamma entrainment therapies, such as GENUS that aim to restore the oscillatory dynamics these interneurons normally support [43].

These results also inform ongoing debates about *top-down* versus *bottom-up* processing deficits in schizophrenia, which are particularly relevant for predictive coding models of schizophrenia [27,44]. Under this framework, delusions and hallucinations may arise from an over-reliance on *top-down* internal priors or from imprecise *bottom-up* sensory prediction errors that fail to update internal models [44]. Planned contrasts revealed preferential involvement of *bottom-up* input layer 4 versus *top-down* input layer 1 and across all *bottom-up* versus *top-down* layers, whereas *top-down* versus *bottom-up* output layers showed no significant difference. A comparable pattern was evident across cell-types, with one informative exception: PV interneurons, which drive feedforward gamma-band inhibition, were more affected than CR and all non-PV subtypes combined, whereas PV and SST did not differ. Rather than a global or uniform breakdown across all circuits, these findings suggest that *bottom-up* signaling deficits are particularly affected. Because SST interneurons modulate both *bottom-up* gamma oscillations [45] and *top-down* information gating [23,24], their joint disruption with PV interneurons may alter precision weighting across microcircuits. However, while brain area significantly predicted effect sizes, deficits did not scale with hierarchical level, suggesting that the primary aberrance reflects a locally conserved microcircuit failure rather than a strictly hierarchical neuropathology.

### Limitations

First, schizophrenia is a clinically heterogeneous disorder. It should be acknowledged that diagnostic criteria spanned multiple historical definitions, and diagnostic-criteria were not readily available in most studies. Second, unequal sampling across brain regions, cell-types, and cortical layers limits statistical power for some analyses. Results based on few studies are less reliable, but we believe they still offer useful, hypothesis-generating information. Third, a high degree of methodological heterogeneity, low sample sizes, and biological sample overlap (driven by the field’s reliance on shared tissue repositories) exists across the published literature. We addressed this by including tissue repository, study, author, sample size, and other ‘nuisance’ covariates in our models. This identified a significant effect of small-sample bias, but this could not explain any of our main results, and no other nuisance factors were significant. Future work should aim to report more complete demographic and technical factors, as well as expand sample size and more broadly sample brain areas, cell types, and cortical layers.

Fourth, these results cannot definitively determine whether interneuron deficits represent a primary phenomenon or arise secondary to impairments. Future research would benefit from exploring the progression of GABAergic deficits in schizophrenia. Similarly, these post-mortem findings do not fully capture dynamic in-vivo circuit states. Importantly, a change in somal labeling within a specific layer localizes the affected source population rather than indicating synaptic pathology [16,46], since interneurons often project beyond their home layer [13,47]. Consequently, these findings map the somal and transcriptional substrates of GABAergic pathology across cortical architecture but not synaptic targets.

Finally, because immunohistochemical detection depends on marker abundance, reduced expression will render some neurons undetectable at conventional resolution and therefore may produce apparent reductions in density that do not reflect absent cells. Consistent with this, Enwright et al. found lower PV-protein per cell body without a deficit in PV neuron number when assessed at high microscope resolution, but at lower resolution, this instead reflected reduced PV neuron number [29]. Similarly, Dienel et al. reported lower SST and PV gene expression without missing neurons [5]. We therefore interpret density as an index of GABAergic pathology that may reflect either absent cells or diminished marker expression within intact cells, and this should be explored further in future work with more sensitive anatomical methods.

### Additional considerations

An additional consideration is that recent work challenges the traditional classification of GABAergic interneurons by PV, CB, and CR [48]. In primate cortex, these markers are largely non-overlapping [49–51] but may also be co-expressed with each other [48] or expressed in non-GABAergic cell types (particularly CB positive neurons), like pyramidal neurons [36,48,52–54]. This is unlikely to affect our study’s conclusions, because the CB studies that explicitly removed CB-stained pyramidal neurons [36,53] arrived at the same conclusions as other studies of CB expression in schizophrenia. In addition, although CB and SST markers can co-express in the same neurons, in macaque V1/V2, the overlap ranges between 36-62%[48]. This helps explain our findings of minimal effect on CB with substantial reductions in SST markers in schizophrenia; these results are likely to be driven by non-overlapping cell-types. This raises the important issue of cellular diversity. All cell-types we have surveyed are genetically diverse and likely have multiple functions. For example, the PV cell class [11] encompasses distinct postsynaptic targets whose relative proportions differ across cortical and subcortical structures. Whether the anatomically similar PV deficits we observe across regions reflect comparable functional consequences therefore remains an open question. Future studies should investigate which specific sub-classes of these PV and SST neurons are affected and what the functional consequence of that is.

## Conclusion

In conclusion, GABAergic interneuron pathology in schizophrenia reflects a pervasive yet microcircuit-specific disruption that preferentially impacts *bottom-up* layers (L2/3/4) and gamma-oscillation associated (PV and SST) components while broadly affecting cortical and subcortical networks. By providing an open-access, denoised resource of EMMs and the underlying standardized dataset, this work enables cell- and layer-specific constraints to be incorporated into circuit models and pharmaceutical interventions. It also offers empirical grounding for resolving how local circuit failures may scale to the symptomatic features of psychosis.

## Data Availability Statement

This review was not registered. However, we make all data and code publicly available (https://github.com/BastosLab/sim-map). This resource includes all code and raw data, as well as pre-processed data including Hedges’ *g* effect sizes for each study and standardized EMMs cell-types, layers, and areas. Other data not included in this resource, Article, or Supplementary Materials can be requested by contacting the corresponding author.

## Acknowledgements

We thank Jadyn Collado for assistance with data collection as well as Helena Mastek and Michelle Franka for the literature research. We also thank Dr. Christine Konradi for her comments to an early version of the manuscript.

## Author contribution statement

A.G.M. and K.M.G. contributed equally to this work. P.J.U. and A.M.B. jointly supervised this work. A.G.M., K.M.G. and A.M.B. conceptualized the study. A.G.M. and K.M.G. performed data collection and analysis and wrote the original draft. All authors contributed to reviewing and editing the manuscript. P.J.U. and A.M.B. acquired funding.

