## Supplemental Information for "GABAergic interneuron pathology in schizophrenia: a systematic review and meta-analysis across cell-types, brain areas, and cortical layers"

Dr. André M. Bastos

Vanderbilt University

301 Wilson Hall

111 21^st^ Ave S

Nashville TN 37203

### PRISMA Checklist

| **Section and Topic** | **Item #** | **Checklist item** | **Location where item is reported** |
| --- | --- | --- | --- |
| TITLE | | |  |
| Title | 1 | Identify the report as a systematic review. | Page 1 |
| ABSTRACT | | |  |
| Abstract | 2 | See the PRISMA 2020 for Abstracts checklist. | Pages 2-3 |
| INTRODUCTION | | |  |
| Rationale | 3 | Describe the rationale for the review in the context of existing knowledge. | Pages 4-5 |
| Objectives | 4 | Provide an explicit statement of the objective(s) or question(s) the review addresses. | Page 5 |
| METHODS | | |  |
| Eligibility criteria | 5 | Specify the inclusion and exclusion criteria for the review and how studies were grouped for the syntheses. | Page 6 |
| Information sources | 6 | Specify all databases, registers, websites, organizations, reference lists and other sources searched or consulted to identify studies. Specify the date when each source was last searched or consulted. | Pages 5-6 |
| Search strategy | 7 | Present the full search strategies for all databases, registers and websites, including any filters and limits used. | Pages 5-6, Supplementary Methods |
| Selection process | 8 | Specify the methods used to decide whether a study met the inclusion criteria of the review, including how many reviewers screened each record and each report retrieved, whether they worked independently, and if applicable, details of automation tools used in the process. | Page 5 |
| Data collection process | 9 | Specify the methods used to collect data from reports, including how many reviewers collected data from each report, whether they worked independently, any processes for obtaining or confirming data from study investigators, and if applicable, details of automation tools used in the process. | Page 6 |
| Data items | 10a | List and define all outcomes for which data were sought. Specify whether all results that were compatible with each outcome domain in each study were sought (e.g. for all measures, time points, analyses), and if not, the methods used to decide which results to collect. | Page 6 |
|  | 10b | List and define all other variables for which data were sought (e.g. participant and intervention characteristics, funding sources). Describe any assumptions made about any missing or unclear information. | Pages 6, 8 |

| Study risk of bias assessment | 11 | Specify the methods used to assess risk of bias in the included studies, including details of the tool(s) used, how many reviewers assessed each study and whether they worked independently, and if applicable, details of automation tools used in the process. | Pages 8-9, Supplementary Table 2 |
| --- | --- | --- | --- |
| Effect measures | 12 | Specify for each outcome the effect measure(s) (e.g. risk ratio, mean difference) used in the synthesis or presentation of results. | Pages 7, 10 |
| Synthesis methods | 13a | Describe the processes used to decide which studies were eligible for each synthesis (e.g. tabulating the study intervention characteristics and comparing against the planned groups for each synthesis (item #5)). | Page 9 |
|  | 13b | Describe any methods required to prepare the data for presentation or synthesis, such as handling of missing summary statistics, or data conversions. | Page 9 |
|  | 13c | Describe any methods used to tabulate or visually display results of individual studies and syntheses. | Pages 9-10, Figure 1A |
|  | 13d | Describe any methods used to synthesize results and provide a rationale for the choice(s). If meta-analysis was performed, describe the model(s), method(s) to identify the presence and extent of statistical heterogeneity, and software package(s) used. | Pages 7, 9-10 |
|  | 13e | Describe any methods used to explore possible causes of heterogeneity among study results (e.g. subgroup analysis, meta-regression). | Page 8 |
|  | 13f | Describe any sensitivity analyses conducted to assess robustness of the synthesized results. | Page 8 |
| Reporting bias assessment | 14 | Describe any methods used to assess risk of bias due to missing results in a synthesis (arising from reporting biases). | Page 8 |
| Certainty assessment | 15 | Describe any methods used to assess certainty (or confidence) in the body of evidence for an outcome. | Pages 8-9 |
| RESULTS | | |  |
| Study selection | 16a | Describe the results of the search and selection process, from the number of records identified in the search to the number of studies included in the review, ideally using a flow diagram. | Page 11, Figure 1A |
|  | 16b | Cite studies that might appear to meet the inclusion criteria, but which were excluded, and explain why they were excluded. | Page 11, Figure 1A |
| Study characteristics | 17 | Cite each included study and present its characteristics. | Pages 11-12, Table 1, Supplementary Table 1 |
| Risk of bias in studies | 18 | Present assessments of risk of bias for each included study. | Pages 11-12, Supplementary Table 2 |
| Results of individual studies | 19 | For all outcomes, present, for each study: (a) summary statistics for each group (where appropriate) and (b) an effect estimate and its precision (e.g. confidence/credible interval), ideally using structured tables or plots. | Figures 2-4, Supplementary Table 1 |
| Results of syntheses | 20a | For each synthesis, briefly summarize the characteristics and risk of bias among contributing studies. | Pages 11-12 |
|  | 20b | Present results of all statistical syntheses conducted. If meta-analysis was done, present for each the summary estimate and its precision (e.g. confidence/credible interval) and measures of statistical heterogeneity. If comparing groups, describe the direction of the effect. | Pages 12-16, Figures 2-4 |
|  | 20c | Present results of all investigations of possible causes of heterogeneity among study results. | Pages 12-16 |
|  | 20d | Present results of all sensitivity analyses conducted to assess the robustness of the synthesized results. | Pages 12-13 |
| Reporting biases | 21 | Present assessments of risk of bias due to missing results (arising from reporting biases) for each synthesis assessed. | Page 12 |
| Certainty of evidence | 22 | Present assessments of certainty (or confidence) in the body of evidence for each outcome assessed. | Pages 12-16 |
| DISCUSSION | | |  |
| Discussion | 23a | Provide a general interpretation of the results in the context of other evidence. | Pages 16-17 |
|  | 23b | Discuss any limitations of the evidence included in the review. | Pages 17-19 |
|  | 23c | Discuss any limitations of the review processes used. | Pages 17-19 |
|  | 23d | Discuss implications of the results for practice, policy, and future research. | Pages 16-19 |
| OTHER INFORMATION | | |  |
| Registration and protocol | 24a | Provide registration information for the review, including register name and registration number, or state that the review was not registered. | Page 20 |
|  | 24b | Indicate where the review protocol can be accessed, or state that a protocol was not prepared. | Page 19 |
|  | 24c | Describe and explain any amendments to information provided at registration or in the protocol. | N/A |
| Support | 25 | Describe sources of financial or non-financial support for the review, and the role of the funders or sponsors in the review. | Page 20 |
| Competing interests | 26 | Declare any competing interests of review authors. | Page 21 |
| Availability of data, code and other materials | 27 | Report which of the following are publicly available and where they can be found: template data collection forms; data extracted from included studies; data used for all analyses; analytic code; any other materials used in the review. | Page 20 |

From: Page, et al. [1]

### Supplementary Methods

#### Additional details on search strategy and selection criteria

The following search terms were used from database inception up to and including Nov 24, 2025: (Schizophrenia OR "Schizophrenia Spectrum and Other Psychotic Disorders"[MeSH] OR SZ) AND (Interneuron* OR "Interneuron density" OR Parvalbumin OR Calbindin OR Calretinin OR Somatostatin OR PV OR CB OR CR OR SST) AND (Immunohistochemistry OR "mRNA expression" OR "gene expression"). Studies were excluded at full-text screening for the following reasons: (1) non-human samples; (2) no cell-specific data; (3) no schizophrenia group; (4) data that could not be extracted or that used indirect proxy measures of density or expression. There were no language restrictions for inclusion.

#### Additional details on data extraction and synthesis

##### Data descriptions

We strictly included interneuron density and mRNA studies for PV, CB, CR, and SST interneurons, including one with both [2]. Included mRNA studies quantified mRNA expression using optical density (OD) analysis, in situ hybridization (ISH), fluorescence in situ hybridization (FISH), quantitative real-time polymerase chain reaction (qPCR), or a combination of FISH/ISH and qPCR. We also included both laminar and non-laminar data, with two studies including both data types [2,3]. For one study [3], raw data were provided by the original authors, who shared an institutional affiliation with members of the present research team.

For the purposes of the present study, we differentiate between brain area (e.g. BA17), brain structure (e.g. visual cortex [VCx], which includes BA17 and 18), and group (PFC, non-PFC cortex, hippocampus, and subcortex). Thus, if a study quantified multiple areas within a structure (i.e. CA1 and CA2), an effect size was computed for each area (Fig. 2C) but grouped for structure-level analysis, publication bias, and heterogeneity analysis (Fig. 2A-B; Appendix table 1). For example, if a study quantified cell density in Brodmann areas 17 (Primary visual cortex, V1) and 18 (Secondary visual cortex, V2), an effect size was computed for each of these areas, and BA17 and 18 (VCx) were also grouped. For study-level analyses, this study would have been included once for area VCx. Structuring the data in this way improved power, as each brain area is a unique measurement, while allowing us to also compare brain structures that may otherwise be sparse. Non-independence between such data was controlled for in the linear modeling with the inclusion of study as a random intercept.

##### Data synthesis

Where standard errors of the mean (SEM) or 95% confidence intervals were reported in place of standard deviations (SD), these metrics were mathematically transformed to standard deviations using standard Cochrane criteria prior to calculating standardized effect sizes. The following equations were used, where 1.96 is for a 95% CI [4]:

$SD = SEM\times\surd n$

$$SD=\frac{CI\times\sqrt{n}}{2\times1.96}$$

For studies reporting data in n≥4 layers, we used the arithmetic mean of layer variances to get a single SD per study (i.e., pooled variance). We then took the arithmetic mean of the layer values (e.g., density or mRNA expression) to get a single mean per study, which we used to compute the average effect size across layers. The following equations were used, $SD_{P}$ represents the pooled standard deviation and $k$ represents the number of layers:

$$SD_{P}=\sqrt{\frac{\sum_{i=1}^{k} \left( n-1 \right){SD}_{i}^{2}}{k(n-1)}}$$

In the linear mixed-effects modelling (LMM), we included subject age, post-mortem interval (PMI), and brain pH. For age, PMI, and pH, we computed the difference between the control and schizophrenia groups; for sex, we computed the ratio of male-to-female subjects for both groups and then the difference between the group’s ratios. We did not treat cohort-paired and cohort-unpaired data differently, because we structured the data in such a way that properly paired cohorts would result in a mathematical difference between the control and schizophrenia groups of zero.

##### False Discovery Rate (FDR) correction

We corrected for multiple comparisons using Benjamini-Hochberg FDR [5]. FDR corrections were performed within hypothesis groups and within comparison groups. This included, for example, across all layers, cell types, and areas for the cortical layer hypothesis. Then, the *p-*values from the Cohen’s *d* comparisons were FDR-corrected separately. This allowed us to test which cell types, layers, and areas were significantly affected in schizophrenia, while correcting for multiple statistical tests run. LMM *p-*values (i.e., significance derived from the model itself) were not FDR-corrected because we ran three, independent models. Subsequent ANOVA, EMM, and dEMM *p-*values were FDR-corrected.

##### mRNA technique analysis

We performed a regression on the mRNA technique subtypes (e.g. FISH/ISH, qPCR, optical density [OD]). An ANOVA did not reveal any significant differences between these methods in the prediction of effect size (F(3,37)=0.49, *p*=0.69). These results suggest that quantification techniques do not impact the resulting effect size when using mRNA.

##### Data visualization

To visualize brain areas included (Figures 1B, 3K), we map our cortical results onto the Desikan-Killiany Atlas [6] and our subcortical results onto the ASEG atlas [7] using the python package yabplot (v0.40, <https://teanijarv.github.io/yabplot/>) [8].

### Demographics of the included studies

#### Supplementary Table 1:‬ Study characteristics‬‬‬‬‬‬‬‬‬‬‬‬‬‬‬‬‬‬‬‬

Papers measuring interneuron density with immunohistochemistry and measuring mRNA expression. The table also includes first author,‬ publication year, and method. Sub-method indicates the specific quantification method used (e.g. neurons/mm‬3‬, mRNA levels [nCi/g]). We also included the‬ medication status, sex, mean age in years, brain pH, post-mortem interval (PMI), and RNA‬ integrity number (RIN) for both healthy control (HC) and schizophrenia (SZ) groups. The tissue‬ repository column indicates the tissue bank(s) from which each study collected samples (if‬ reported). The reagent manufacturer for the antibody/reagent used is included for each cell type. Medication status and RIN had insufficient data to be included in any model. We also included from which figure or table we extracted the data, and the method of data extraction (e.g. direct from a table or WebPlotDigitizer manual extraction).‬‬‬‬‬‬‬‬‬‬‬‬‬‬‬ We include the resulting effect size (Hedges’ g) ± variance of the effect size for each cell type, method, or cortical layer. In cases where laminar data is reported and reported across at least 4 layers, we also include the mean effect size ± pooled variance across layers. Also included is the overall risk of bias with the main reasons for this appraisal (see Supplementary Table 2).‬‬‬‬‬‬‬‬‬‬‬‬

‬‬‬‬‬‬‬‬‬‬‬‬

### Supplementary Analyses

##### Cell type-method interaction analysis

We performed a regression on the interaction between cell type and method (e.g. IHC/mRNA). Model 1 (demographics and technical regression model) revealed that the effect size of IHC (EMM=–0.75) was nominally larger than mRNA (EMM=–0.33), after controlling for various technical factors, cell type, and brain area. Furthermore, multivariate adjustment in the interaction LMM model revealed that methodological differences depend significantly on the specific cell type investigated (*F*(3, 121)=3.26, *p*=0.024). Specifically, the main effect of method (β=–0.15, SE=0.049, *t*(121)=–3.16, *p*=0.0020), suggesting that IHC deficits may be roughly 0.31 standard deviations larger overall than mRNA when accounting for cell-type interactions.

Cell-types exhibited largely similar patterns between methods, with the largest differences emerging from SST interneurons. PV interneurons were more affected in mRNA expression (EMM=–0.66, 95% CI [–0.83, –0.49], *p*<0.0001) than IHC (EMM=–0.58, 95% CI [–0.77, –0.37], *p*<0.0001). CB interneuron density (IHC) was reduced in schizophrenia compared to controls (EMM=–0.30, 95% CI [–0.60, –0.00040], *p*=0.0497), whereas CB mRNA expression was increased in schizophrenia compared to controls (EMM=0.39, 95% CI [0.034, 0.74], *p*=0.032). CR interneuron density (EMM=–0.17, 95% CI [–0.44, 0.097], *p*=0.21) and mRNA expression (EMM=0.071, 95% CI [–0.21, 0.35], *p*=0.62) were not affected in schizophrenia. Finally, SST interneuron density (EMM=–1.064, 95% CI [–1.422, –0.710], *p*<0.0001) was substantially more affected than SST mRNA expression (EMM=–0.68, 95% CI [–0.86, –0.51], *p*<0.0001), though both were significantly reduced.

##### Cell type-brain area analysis

We performed an additional regression to see how cell types changed in each area. An interaction model was not used here due to sparsity of areas for some cell types (namely CR interneurons). Method and sample size were included as covariates, along with study and tissue repository as random effects. Generally, CB interneurons were the least affected cell types across areas, whereas SST interneurons had the most negative EMMs. CR interneurons exhibited more modest effects across areas and PV interneurons were slightly less affected than SST.

##### Cell type-cortical layer analysis

We performed an additional regression to see how each cell type changed across cortical layers. Again, an interaction model was not used here for the same reasons mentioned above. Sample size as included as a covariate, along with study and tissue repository as random effects. However, method had to be removed due to sparsity of cell types (CB and SST) containing sufficient data from multiple layers across both methods and both random effects.

##### Reagent manufacturer analysis

We also extracted the reagent manufacturer per cell type when reported by study. We performed an additional regression to see if the manufacturer of the reagent for each cell type significantly predicted effect size. In this model, we also included sample size, ∆pH, ∆PMI, ∆Age, and cell type as additional factors in the model, as well as random intercepts for author and tissue repository. Consistent with other models, cell type and sample size were significant predictors of effect size (*p*<0.0001). Reagent manufacturer did not significantly predict effect size (*F*(7, 41)=2.086, *p*=0.065). Other reagent information was inconsistently reported, restricting analyses on other factors.

#### Supplementary Table 2: Risk of bias table

| **Study** | **Participant Matching** | **Covariate Control** | **Medication Confounding** | **Sample Size** | **Data Quality** | **Overall** | **Primary Justification for High-Risk Designation** |
| --- | --- | --- | --- | --- | --- | --- | --- |
| Sakai *et al.,* 2007 (IHC) | 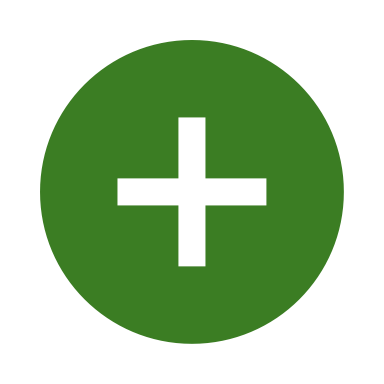 | 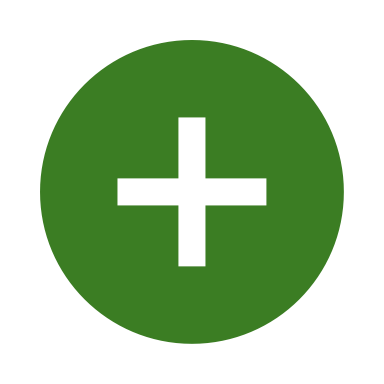 | 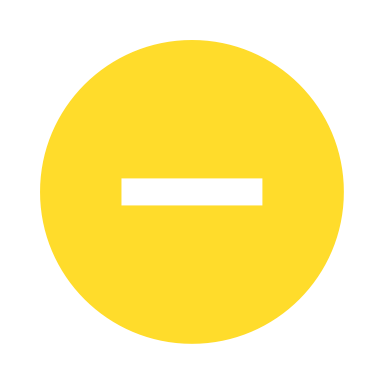 | 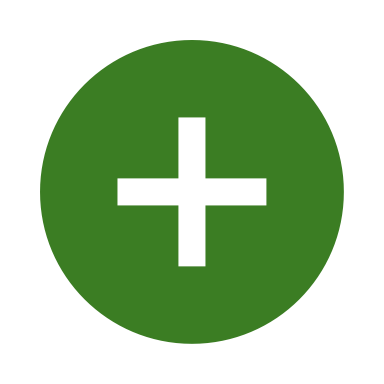 | 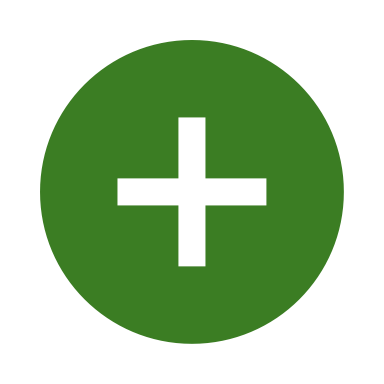 | 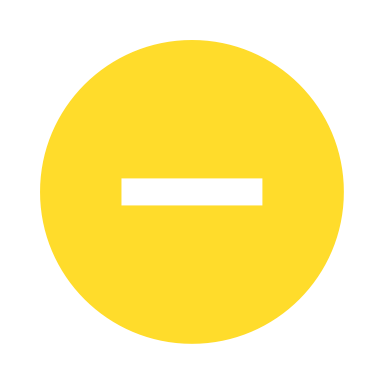 |  |
| Tooney & Chahl 2004 (IHC) | 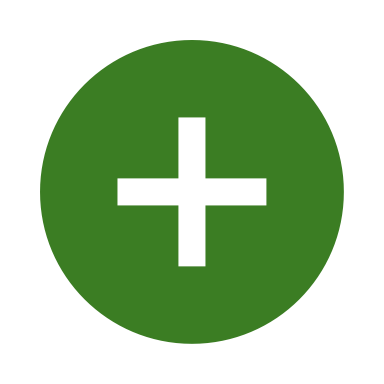 | 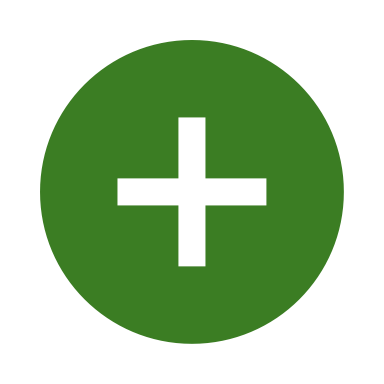 | 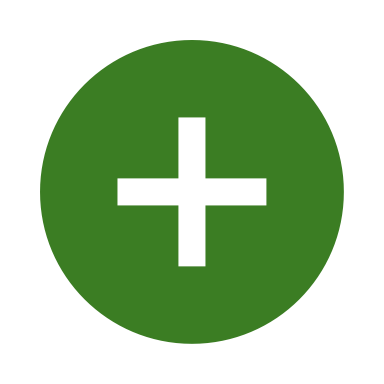 | 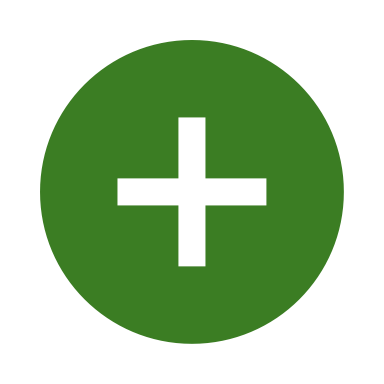 | 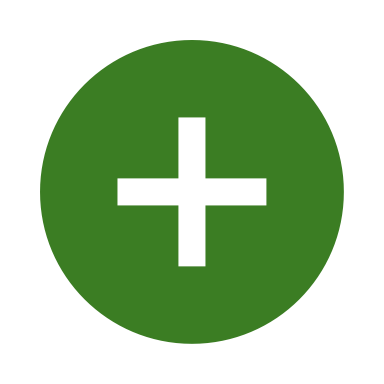 | 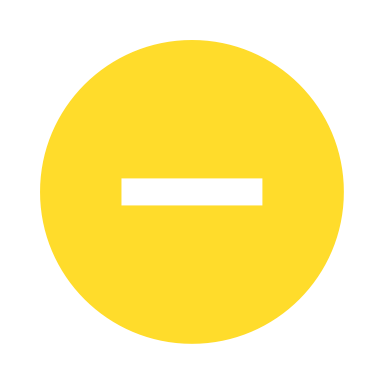 |  |
| Reynolds *et al.,* 2002 (IHC) | 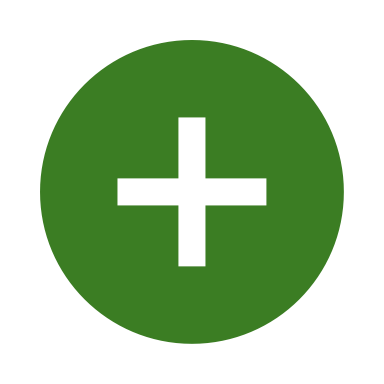 | 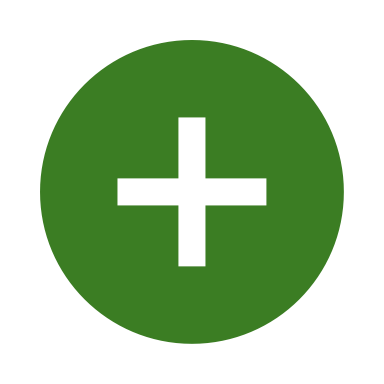 | 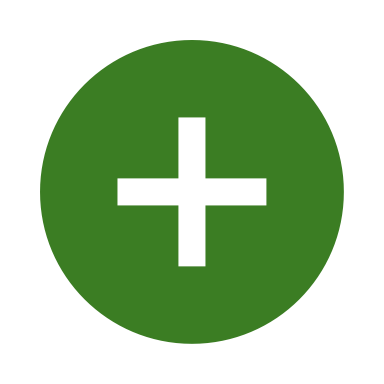 | 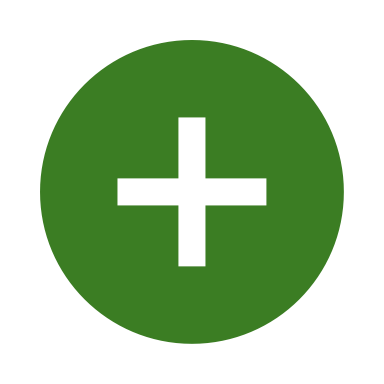 | 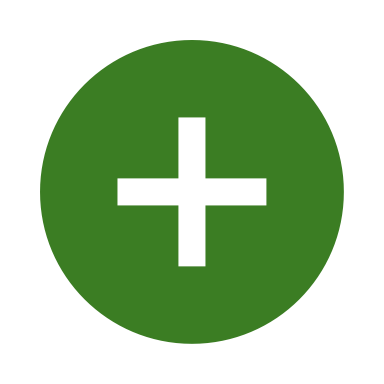 | 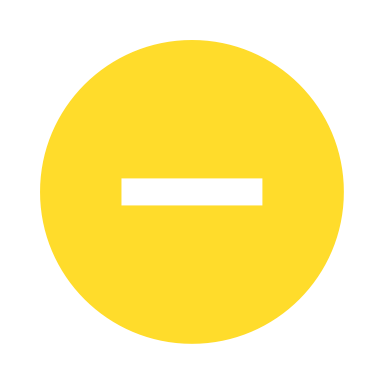 |  |
| Beasley & Reynolds, 1997 (IHC) | 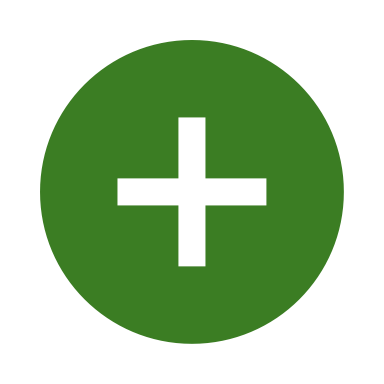 | 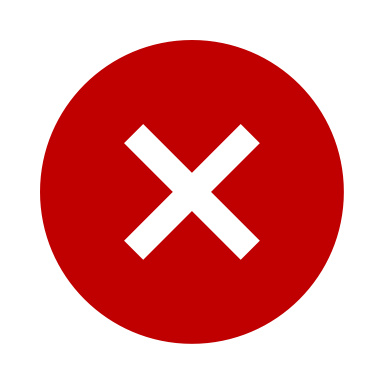 | 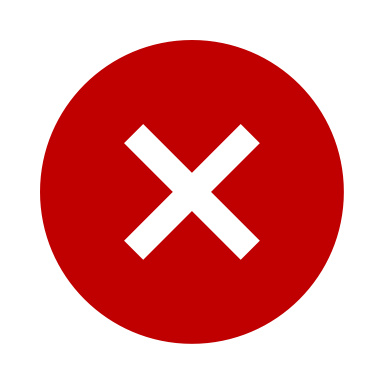 | 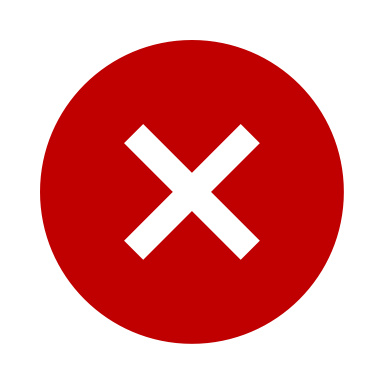 | 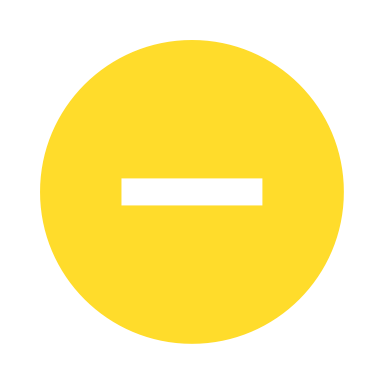 | 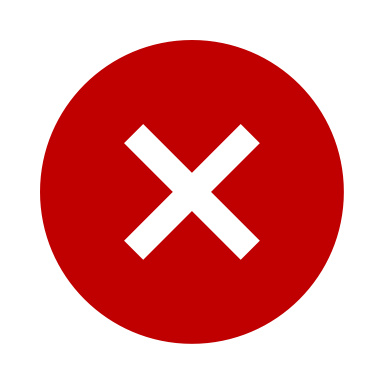 | Absence of baseline tissue stability and brain pH verification records |
| Beasley *et al.,* 2002 (IHC) | 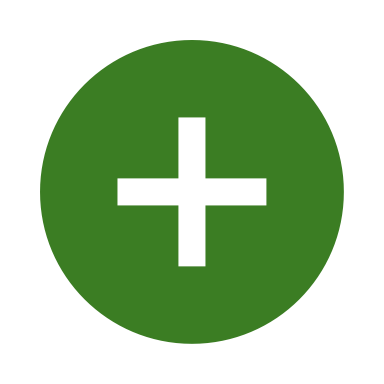 | 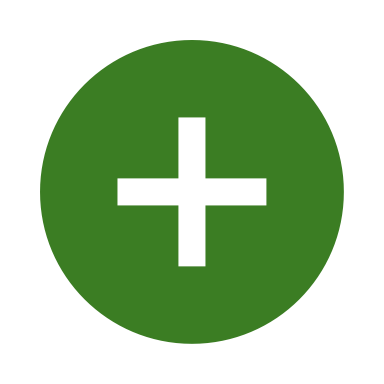 | 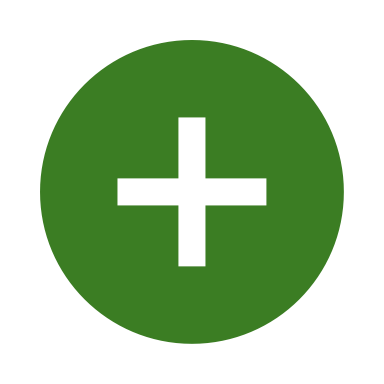 | 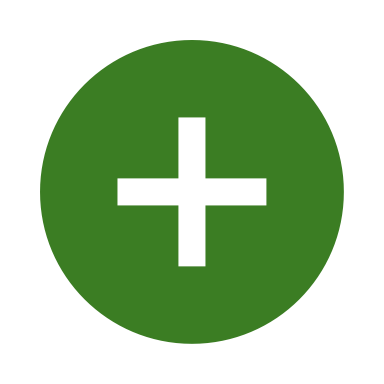 | 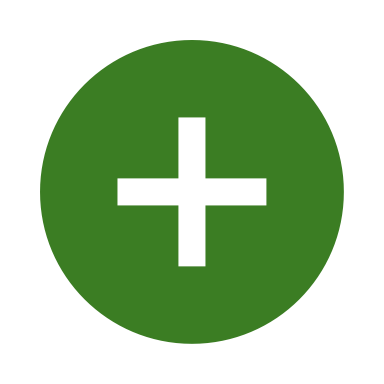 | 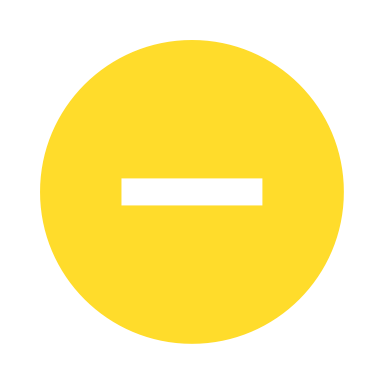 |  |
| Reynolds & Beasley, 2001 (IHC) | 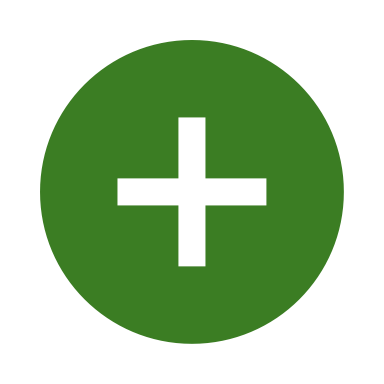 | 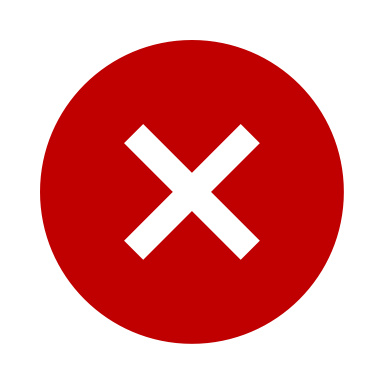 | 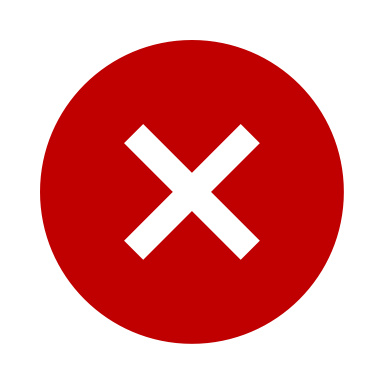 | 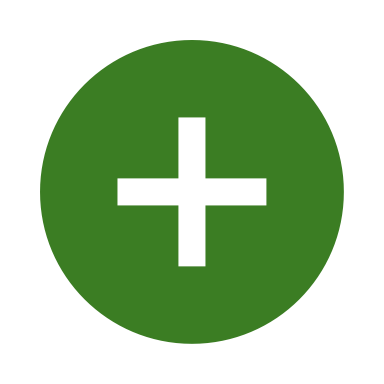 | 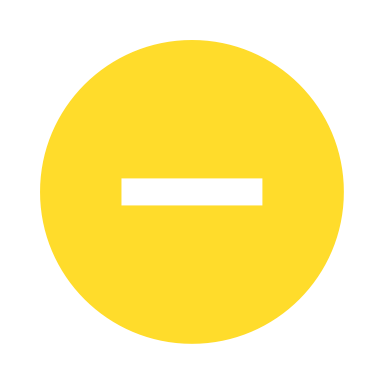 | 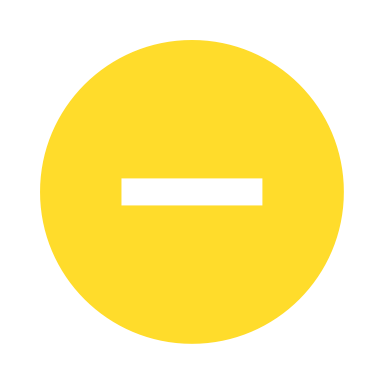 |  |
| Enwright *et al.,* 2016 (IHC) | 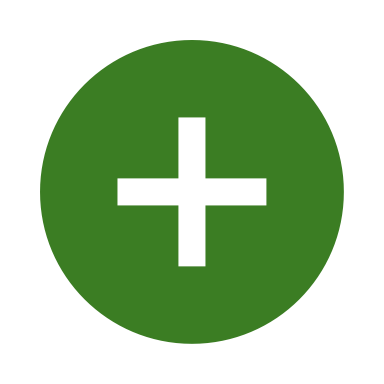 | 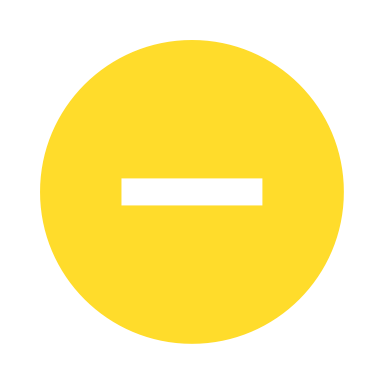 | 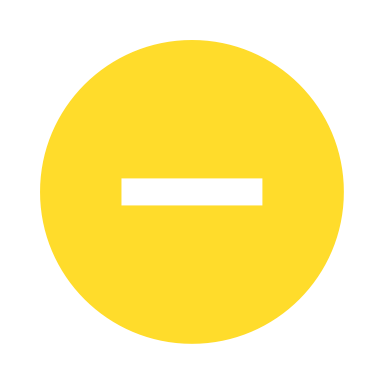 | 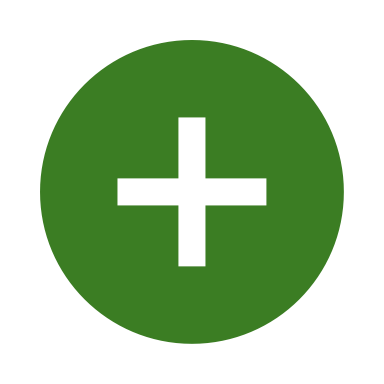 | 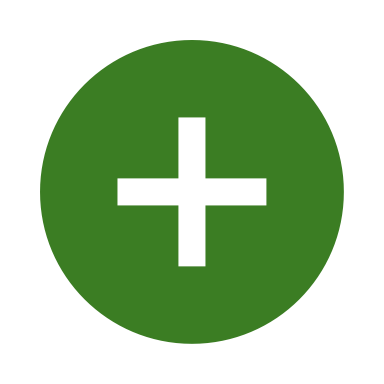 | 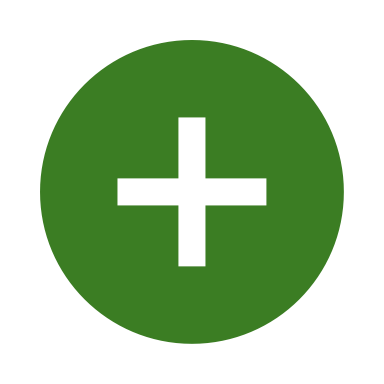 |  |
| Batiuk *et al.,* 2022 (IHC) | 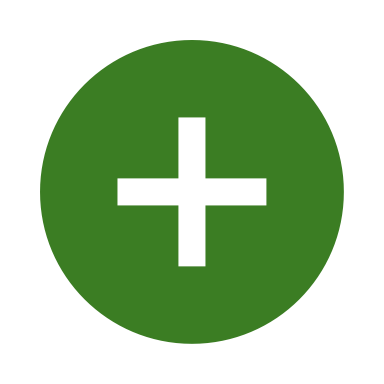 | 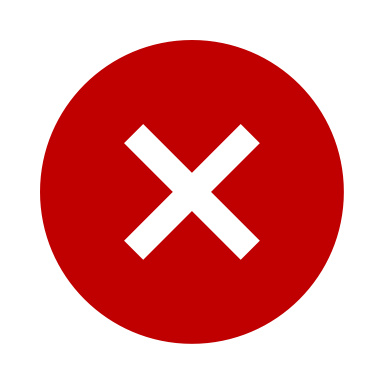 | 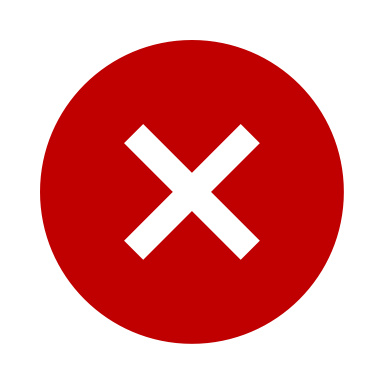 | 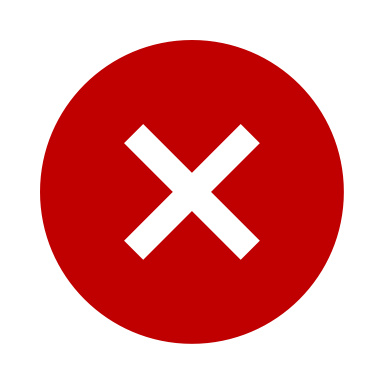 | 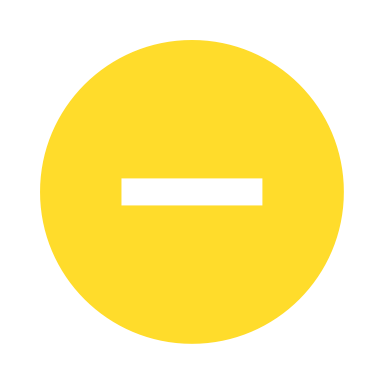 | 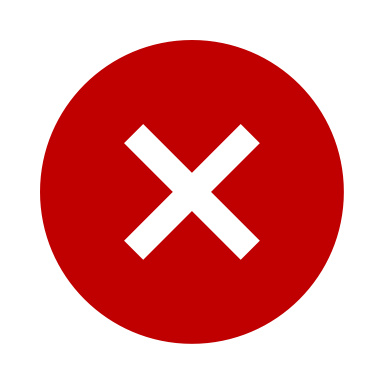 | Blended 4 international repositories without reporting site-specific pH/variances. |
| Pantazopoulos *et al.,* 2007 (IHC) | 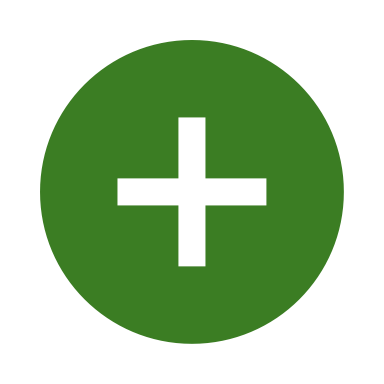 | 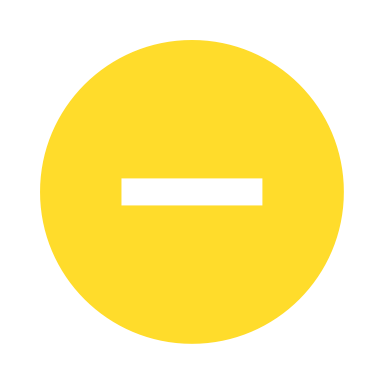 | 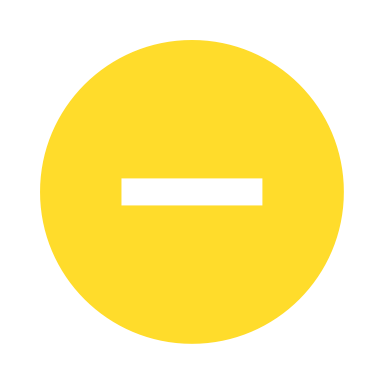 | 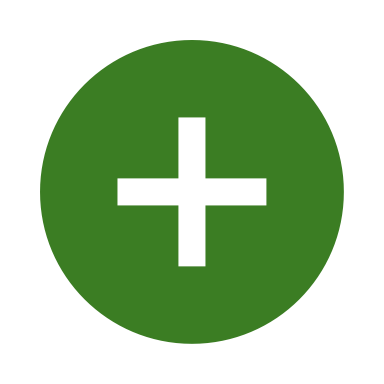 | 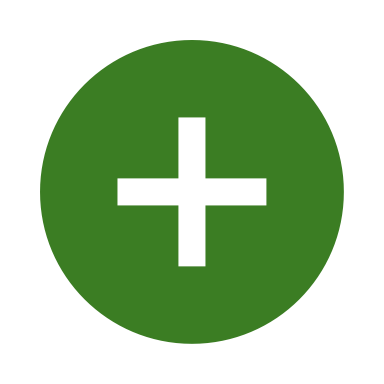 | 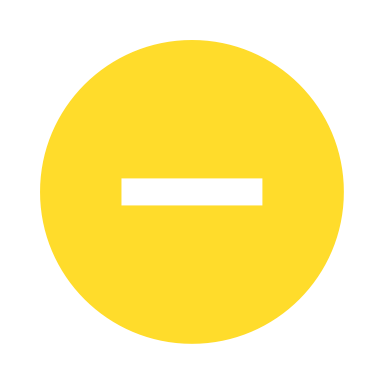 |  |
| Woo *et al.,* 1997 (IHC) | 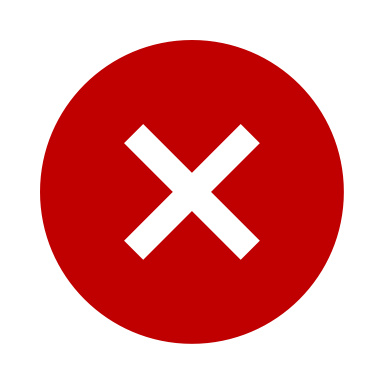 | 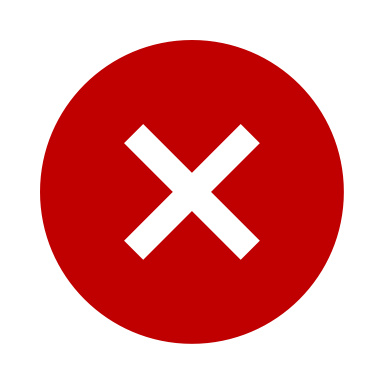 | 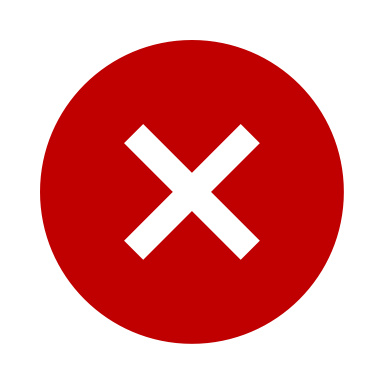 | 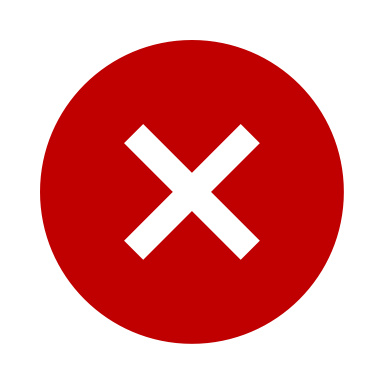 | 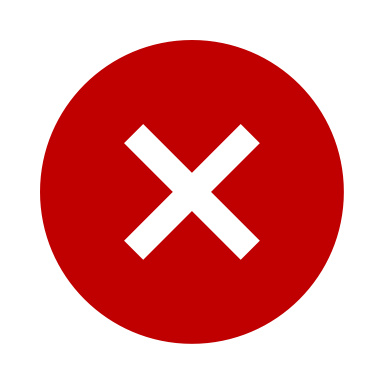 | 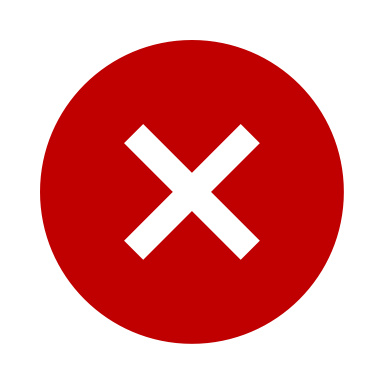 | Historical cohort; missing pH reporting and clear antipsychotic histories. |
| Kalus *et al.,* 1997 (IHC) | 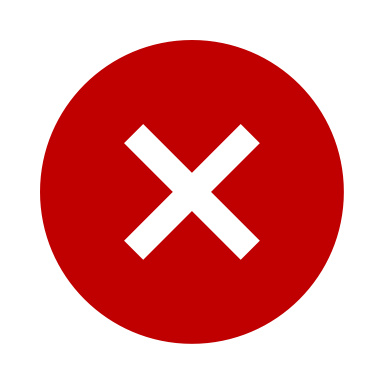 | 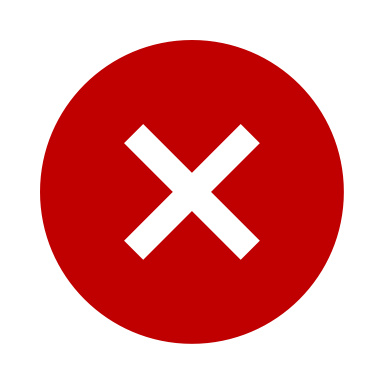 | 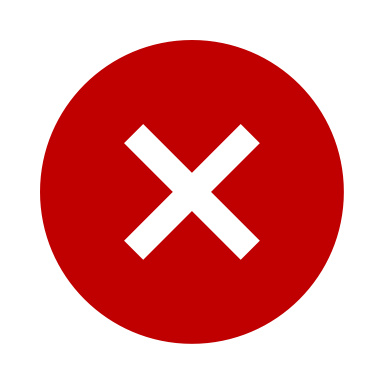 | 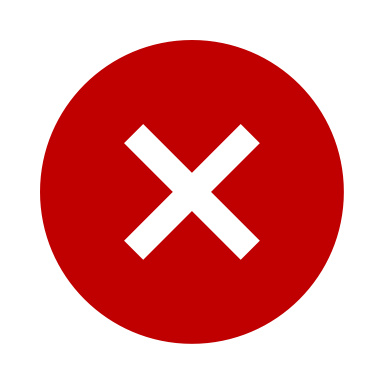 | 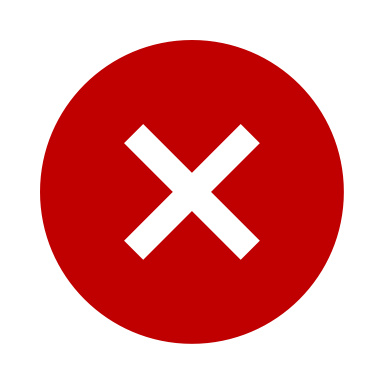 | 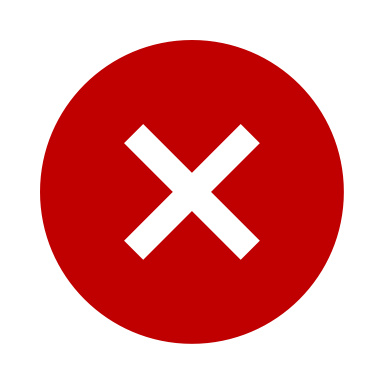 | Small sample size and absence of baseline brain pH data |
| Daviss & Lewis, 1995 (IHC) | 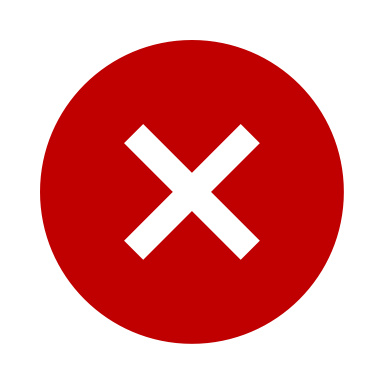 | 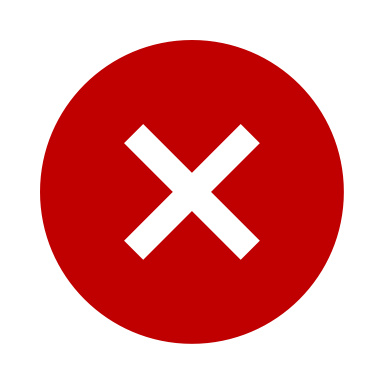 | 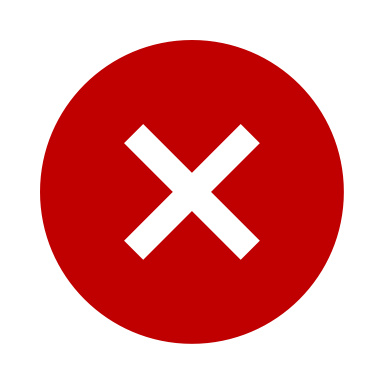 | 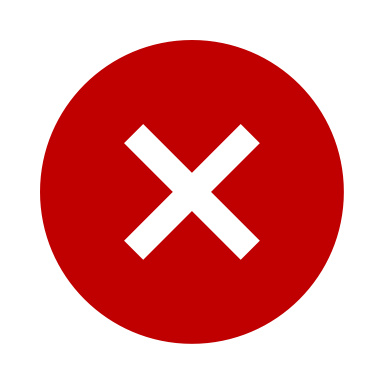 | 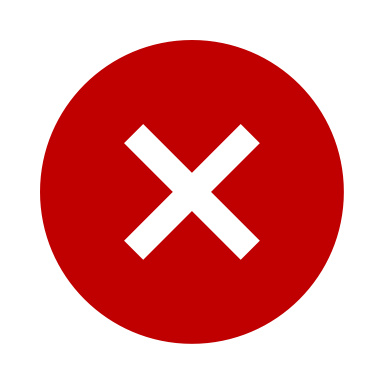 | 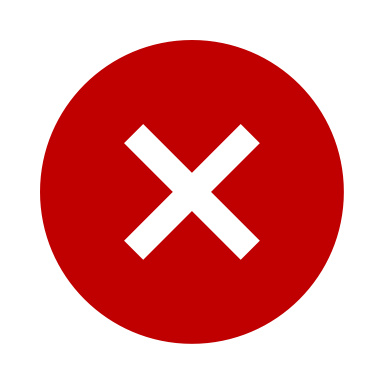 | Historical and small cohort; missing pH, demographic matching, and medication reporting. |
| Dupper, 2013 (IHC) | 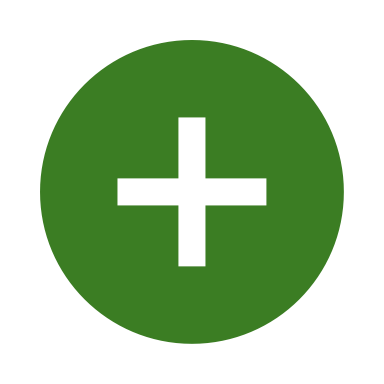 | 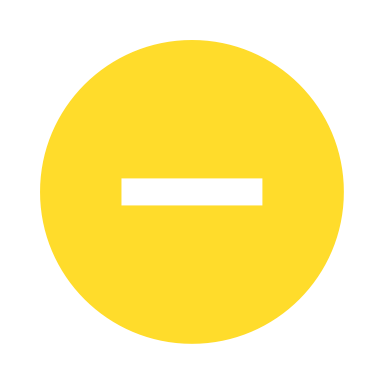 | 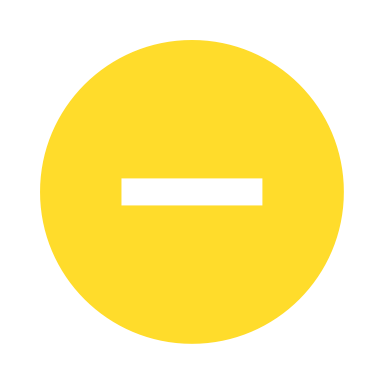 | 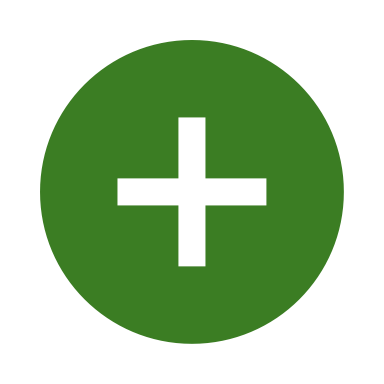 | 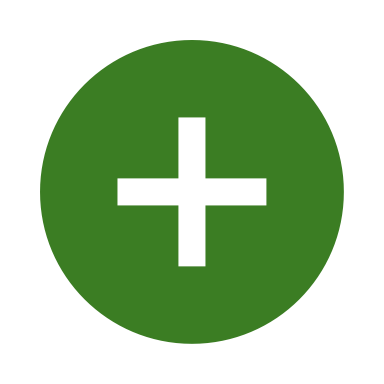 | 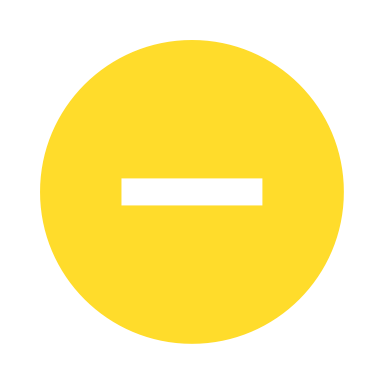 |  |
| Chance *et al.,* 2005 (IHC) | 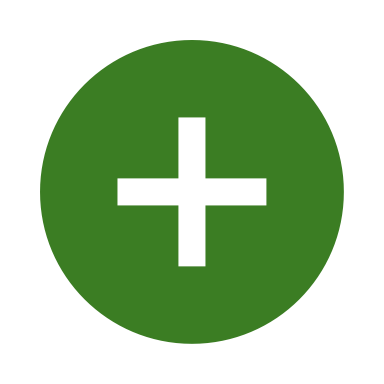 | 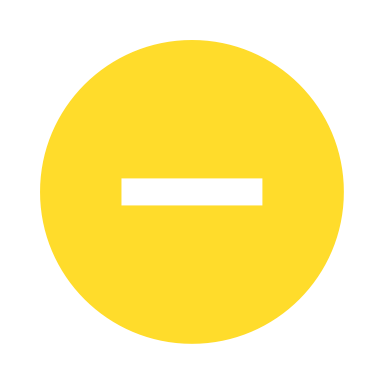 | 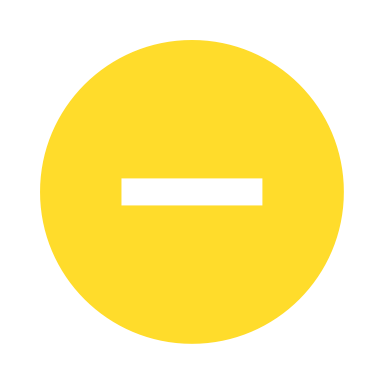 | 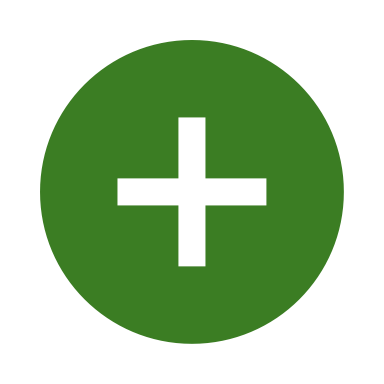 | 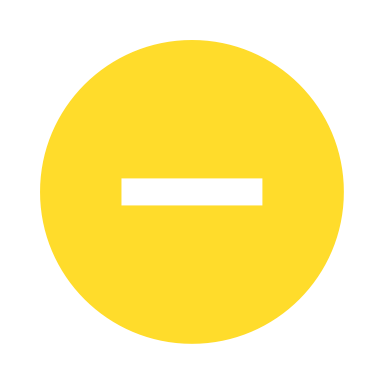 | 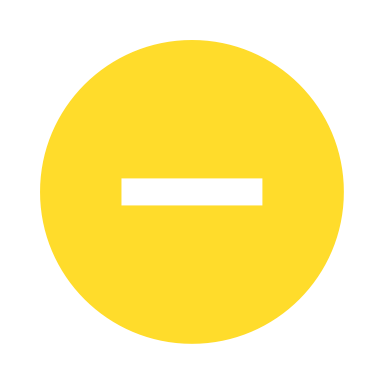 |  |
| Wang *et al.,* 2011 (IHC) | 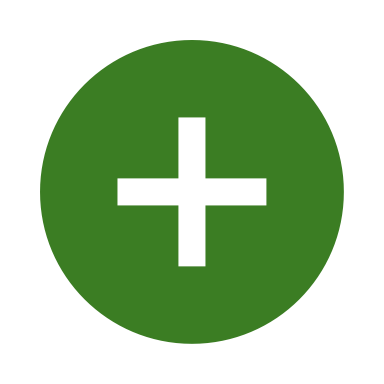 | 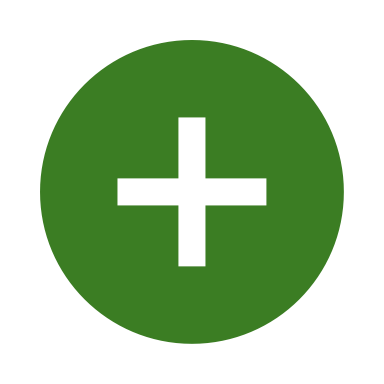 | 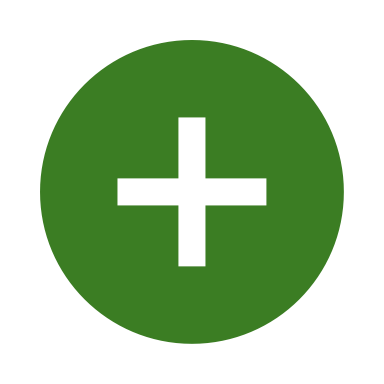 | 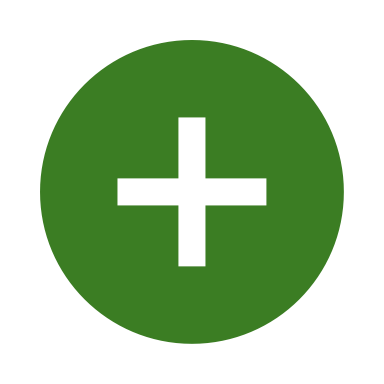 | 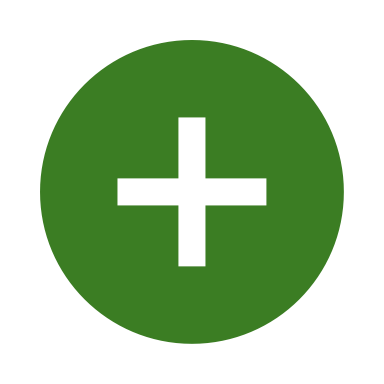 | 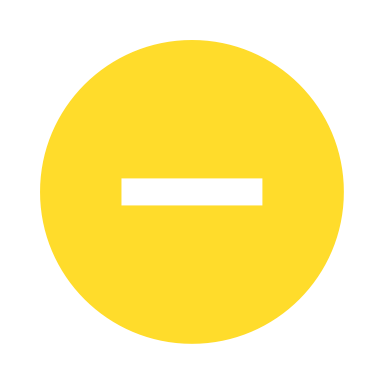 |  |
| Wheeler *et al.,* 2006 (IHC) | 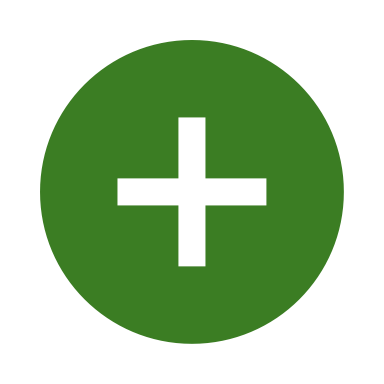 | 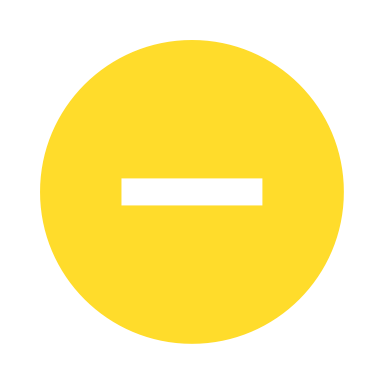 | 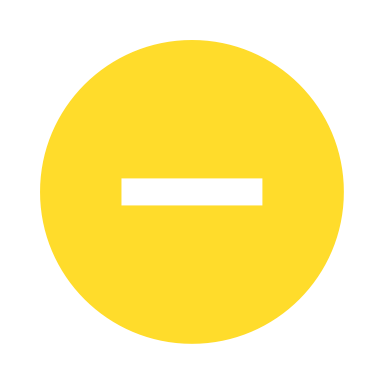 | 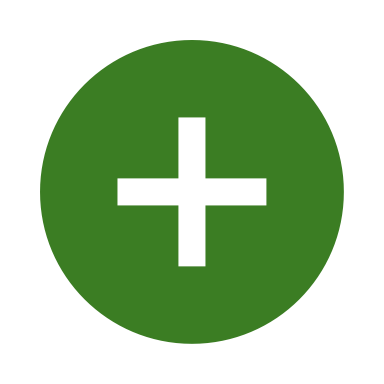 | 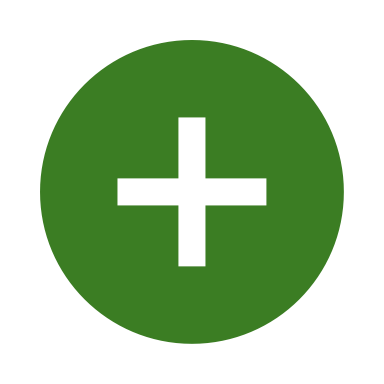 | 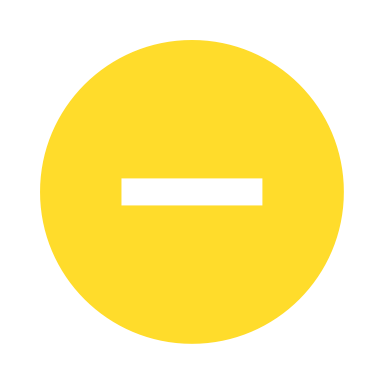 |  |
| Steullet *et al.,* 2018 (IHC) | 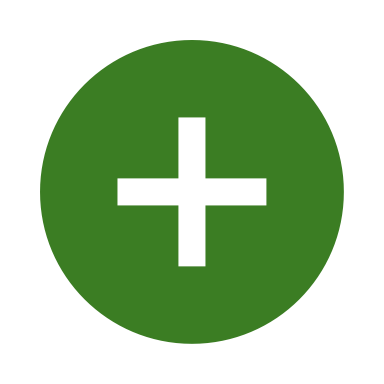 | 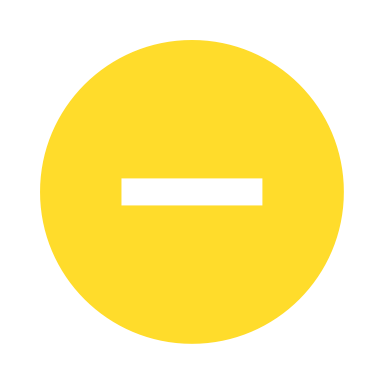 | 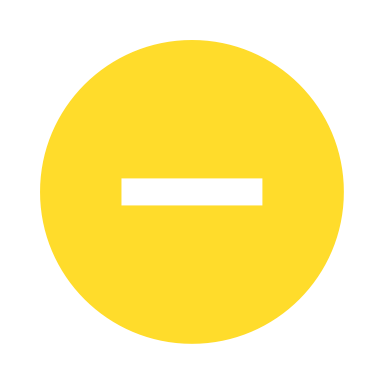 | 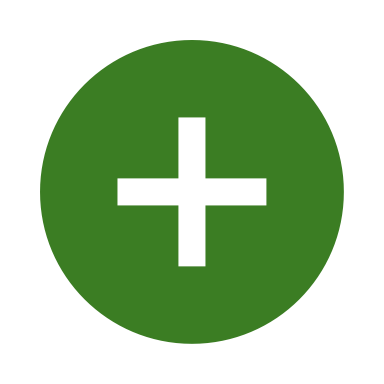 |  |  |  |
| Konradi *et al.,* 2011 (IHC) |  |  |  |  |  |  |  |
| Zhang & Reynolds 2002 (IHC) |  |  |  |  |  |  |  |
| Farmer *et al.,* 2023 (IHC) |  |  |  |  |  |  |  |
| Falkai *et al.,* 2016 (IHC) |  |  |  |  |  |  |  |
| Kilonzo *et al.,* 2020 (IHC) |  |  |  |  |  |  |  |
| Adorjan *et al.,* 2020 (IHC) |  |  |  |  |  |  |  |
| Holt *et al.,* 1999 (IHC) |  |  |  |  |  |  | Historical and small cohort; limited pH and medication reporting |
| Pantazopoulos *et al.,* 2017 (IHC) |  |  |  |  |  |  |  |
| Dienel *et al.,* 2025 (mRNA) |  |  |  |  |  |  |  |
| Dienel *et al.,* 2023 (mRNA) |  |  |  |  |  |  |  |
| Joshi *et al.,* 2015 (mRNA) |  |  |  |  |  |  |  |
| Morris *et al.,* 2008 (mRNA) |  |  |  |  |  |  |  |
| Takahashi *et al.,* 2002 (mRNA) |  |  |  |  |  |  |  |
| Hashimoto *et al.,* 2003 (mRNA) |  |  |  |  |  |  |  |
| Fung *et al.,* 2010 (mRNA) |  |  |  |  |  |  |  |
| Nakatani *et al.,* 2006 (mRNA) |  |  |  |  |  |  |  |
| Volk *et al.,* 2016 (mRNA) |  |  |  |  |  |  |  |
| Purves-Tyson *et al.,* 2021 (mRNA) |  |  |  |  |  |  |  |
| Fung *et al.,* 2014 (mRNA) |  |  |  |  |  |  |  |
| Chung *et al.,* 2018 (mRNA) |  |  |  |  |  |  |  |
| Tsubomoto *et al.,* 2019 (mRNA) |  |  |  |  |  |  |  |
| Volk *et al.,* 2012 (mRNA) |  |  |  |  |  |  |  |
| Woo *et al.,* 2008 (mRNA) |  |  |  |  |  |  |  |
| Konradi *et al.,* 2011 (mRNA) |  |  |  |  |  |  |  |
| Hashimoto *et al.,* 2008 (mRNA) |  |  |  |  |  |  |  |
| Okuda *et al.,* 2024 (mRNA) |  |  |  |  |  |  |  |
| Chung *et al.,* 2015 (mRNA) |  |  |  |  |  |  |  |
| Mellios *et al.,* 2009 (mRNA) |  |  |  |  |  |  |  |

Risk of bias across six risk domains (Medication, Participant Matching/Control, Covariate Confounding, and Sample Size). Green (+) indicates low risk, yellow (-) indicates some concerns/moderate risk, and red (X) indicates high risk. Primary justifications for high-risk designations are provided in the rightmost column and are frequently related to missing brain pH records, baseline tissue stability, and sample size limitations. Data quality refers to the reliability of outcome measurement (quantification method, tissue processing, data extraction transparency, technical confounds, etc.).

#### Supplementary Fig. 1: Publication bias funnel plots for all datasets

Funnel plots for visual inspection of publication bias. Studies are indicated as blue dots and 95% CI represented as inverted triangle shapes in dotted lines. Statistical significance of bias is assessed with Egger’s regression.

#### Supplementary Fig. 2: Estimated marginal means of cell-types and brain areas

Estimated marginal means (EMMs) with 95% CI from Model 3 (layer model) for (a) cell type and (b) area. Grand mean plotted in dashed line, with significance assessed via a Wald F-Test against zero (black asterisk) and the grand mean (red asterisk).

#### Supplementary Fig. 3: Estimated marginal means of brain areas

Estimated marginal means (EMMs) for brain areas from Model 2. 95% CI shown in error bars with significance stars as black asterisks for deviations from 0. Dashed line represents the grand mean, and red asterisks represent significant deviations from this average. Areas are colored based on their hierarchical grouping with grey for outside of hierarchy, blue for posterior, red for anterior, yellow for entorhinal cortex (EC), and purple for hippocampus.

#### Supplementary Fig. 4: Estimated marginal means of brain areas by cell-type

Estimated marginal means (EMMs) from the cell-type brain area analysis. 95% CI shown in error bars with significance stars as black asterisks representing deviation from zero. PV interneurons shown in dark blue, CB interneurons in green, CR interneurons in orange, and SST interneurons in light blue. The “average” effect size across cell types per area from Model 2 is shown in black dot with error bars. Note that this is not the arithmetic mean of the EMMs, but rather the model estimates for brain area when controlling for all cell types.

#### Supplementary Fig. 5: Estimated marginal means of cortical layers by cell-type

Estimated marginal means (EMMs) from the cell-type cortical layer analysis. 95% CI shown in error bars with significance stars as black asterisks representing deviation from zero. PV interneurons shown in dark blue, CB interneurons in green, CR interneurons in orange, and SST interneurons in light blue.

#### Supplementary Table 3: Included studies divided by region and method

| **Region/Group** | **Method** | **n_s_** | **Percentage of Total (%)** |
| --- | --- | --- | --- |
| PFC | IHC | 11 | 25.00% |
|  | mRNA | 15 | 34.09% |
| Cortex (PFC excluded) | IHC | 5 | 11.36% |
|  | mRNA | 3 | 6.82% |
| Hippocampus | IHC | 5 | 11.36% |
|  | mRNA | 1 | 2.27% |
| Subcortex | IHC | 3 | 6.82% |
|  | mRNA | 1 | 2.27% |

Percentage and number of studies divided by the brain region and methodology used. Some studies quantified multiple brain regions/groups and methods.

#### Supplementary Table 4: Heterogeneity testing results

| Method Type | PFC | | | | Hippocampus | | | Cortex | | |
| --- | --- | --- | --- | --- | --- | --- | --- | --- | --- | --- |
|  | *n_o_* | | I^2^ (%) | $\boldsymbol{\tau}^{\mathbf{2}}$ | *n_o_* | I^2^ (%) | $\boldsymbol{\tau}^{\mathbf{2}}$ | *n_o_* | I^2^ (%) | $\boldsymbol{\tau}^{\mathbf{2}}$ |
| PV IHC | 8 | 0.00 | | 0.00 | 5 | 77.97 | 0.48 | 5 | 52.00 | 0.18 |
| CB IHC | 11 | 68.79 | | 0.14 | 1 | – | – | 2 | – | – |
| CR IHC | 7 | 0.00 | | 0.00 | 1 | – | – | 5 | 0.00 | 0.00 |
| SST IHC | 2 | – | | – | 0 | – | – | 1 | – | – |
| PV mRNA | 7 | 0.00 | | 0.00 | 1 | – | – | 1 | – | – |
| CB mRNA | 5 | 29.53 | | 0.024 | 0 | – | – | 0 | – | – |
| CR mRNA | 0 | – | | – | 2 | – | – | 1 | – | – |
| SST mRNA | 11 | 56.99 | | 0.088 | 1 | – | – | 5 | 63.74 | 0.31 |

*n_s_* indicates the number of studies for that method type within the indicated group (PFC, hippocampus, or non-PFC cortex). There was insufficient data to explore heterogeneity in subcortical areas. I^2^ is indicated as a percentage (%) for *n_s_* ≥ 5. $\tau^{2}$ indicated for *n_s_* ≥ 5.

#### Supplementary Table 5: Leave-one-author-out sensitivity analysis

| Parameter | Mean | Standard Deviation | t-statistic | *p*-value |
| --- | --- | --- | --- | --- |
| Model Intercept | -0.690 | 0.381 | -7.471 | *p* < 0.0001 |
| Age | 0.039 | 0.066 | 2.413 | 0.028 |
| PMI | 0.029 | 0.041 | 2.876 | 0.011 |
| Sample size | 0.003 | 0.002 | 6.295 | *p* < 0.0001 |
| PV | -0.722 | 0.078 | -38.163 | *p* < 0.0001 |
| CB | 0.223 | 0.070 | 13.156 | *p* < 0.0001 |
| SST | -0.662 | 0.061 | -44.957 | *p* < 0.0001 |
| Method (mRNA) | 0.420 | 0.468 | 3.706 | 0.002 |

Mean coefficient and standard deviation achieved across leave-one-out iterations. Also, the t-statistic and *p*-value from a *t-*test across leave-one-out iterations is included, where a significant *p*-value indicates stability.

#### Supplementary Table 6: Number of tests per FDR Family

| **Family Description** | **N Tests** |
| --- | --- |
| One-Sample Laminar Effect Sizes | 35 |
| Cohen's Laminar Comparison | 36 |
| One-Sample Cellular Effect Sizes (Within Area) | 18 |
| Cohen's Cell-Cell Comparisons | 15 |
| One-Sample Effect Sizes within PFC/Hipp | 13 |
| Cohen's Comparisons across groups (e.g., PV PFC vs. Cortex) | 22 |
| Demographics (Model 1) Anova | 9 |
| H2/3 (Model 2) Cell/Area Anova | 5 |
| H1 (Model 3) Laminar Anova | 6 |
| H1 (Layers) EMMs: Cell | 4 |
| H1 (Layers) EMMs: Layer | 6 |
| H1 (Layers) EMMs: Area | 6 |
| H2/3 (Area/Cell) EMMs: Cells | 4 |
| H2/3 (Area/Cell) EMMs: Areas | 14 |
| BU/TD Comparisons dEMM Layers | 3 |
| BU/TD Comparisons dEMM Cell/Area | 9 |
| H1 BU/TD EMMs | 6 |
| H2/3 Cell/Area EMMs | 9 |

Number of tests per multiple comparison correction. Families are tests grouped by hypothesis that are corrected together. H=Hypothesis. BU=Bottom-up. TD=Top-down. EMM=estimated marginal mean. dEMM=difference in estimated marginal mean.

#### Supplementary Table 7: Reagent manufacturer extraction information

We provide the extracted reagent manufacturer data in an external table. This included the study information (first author, publication year, method, area, cell type(s)), as well as relevant reagent information. This included reagent type, manufacturer/source, and other technical factors.

8.	Toomas Erik Anijärv, jadenecke, Anthony (Tony) Barrows. teanijarv/yabplot: 0.4.0. 2026.
